# Generational selection, transcriptomics and functional characterization reveal the impact of environmental pollutants on the evolution of insecticide resistance in malaria vectors

**DOI:** 10.64898/2026.03.10.710841

**Authors:** Abdullahi Muhammad, Sulaiman S. Ibrahim, Helen Irving, Talal Al-Yazeedi, Jack Hearn, Mark J.I. Paine, Charles S. Wondji

## Abstract

Insecticide resistance is threatening malaria control. While the evolution and spread of resistance has been linked to scale-up in the distribution of public health insecticides, the role of environmental pollutants such as the polyaromatic hydrocarbons (PAHs) from industrial and agricultural use remains largely uncharacterized. The PAHs are potent ligands of the aryl hydrocarbon receptor (Ahr) transcription factors involved in the regulation of xenobiotic metabolizing enzymes, and potentially involved in insecticide resistance. Here, using field insecticide-resistant (Auyo) *An. coluzzii* and a laboratory-susceptible colony (Ngousso), we conducted a multi-generational selection experiment using naphthalene, fluorene and a mixture of both PAHs. After ten generations, the changes in susceptibility to insecticides were monitored using WHO bioassays and whole-transcriptome analysis (RNASeq) was conducted. Compared with the non-selected colony lines, PAH exposures significantly reduced pyrethroid and DDT resistance in the field population, suggesting fitness cost associated with established resistance. In contrast, Ngousso showed a significant increase in DDT resistance (*p* = 0.01) at the tenth generation. A significant increase in permethrin resistance was also observed at the seventh generation (p = *0.03*). Several candidate genes from the major detoxification classes were overexpressed in the selected lines (including GSTe2, CYP6Z1, and CYP6P4); the most consistent were CYP6M4 and CYP4C27, as well as those from the Ahr pathway. Heterologous expression of *CYP6M4* revealed its ability to metabolise pyrethroids, including permethrin, deltamethrin, and α-cypermethrin, as well as PAHs (naphthalene and fluorene). These findings establish the role of environmental pollutants as additional drivers of metabolic insecticide resistance in *An, coluzzii*.

## 1.1 Background

Malaria is a major public health concern in sub-Saharan Africa with its associated morbidity and mortality affecting the most vulnerable in the society (WHO, 2024). Malaria control relies on the use of chemical-based interventions in the form of long-lasting insecticide-treated nets and indoor residual spraying (Bhatt et al., 2015). However, the emergence and escalation of insecticide resistance (Ranson and Lissenden, 2016; Riveron et al., 2018; WHO, 2012) is threatening the success of controlling mosquito vectors of malaria. Increased selection pressures from insecticide based tools used for public health vector control, and the widespread use of agricultural pesticides and from environmental pollutants have been linked to insecticide resistance in malaria vectors (Chouaïbou et al., 2016; Kamdem et al., 2017a; Nkya et al., 2014; Surendran et al., 2019).

Exposure to pollutants has been found to directly affect the insecticide susceptibility of malaria vectors through various mechanisms, including overexpression of detoxification genes implicated in insecticide resistance and/or metabolism. For example, exposure of mosquito larvae to soap and disinfectants led to reduced mortality of adult *An. gambiae* to permethrin (Antonio-Nkondjio et al., 2014). Other environmental pollutants found to impact the insecticide susceptibility of mosquitoes include fluoranthene (Poupardin et al., 2012) in which pre-exposure of *Aedes aegypti* larvae to fluoranthene prior to selection with permethrin showed an increase in insecticide resistance and overexpression of many detoxification genes compared to the non-selected lines. In a related study, the rearing of *An. arabiensis* larvae under high concentrations of lead nitrate, copper nitrate and cadmium chloride led to an increase in the expression of detoxification genes including P450s, and esterases (Oliver and Brooke, 2018). Similar responses were observed with *An. arabiensis* larval exposure to inorganic fertilizer (Samuel et al., 2020), which affected insecticide resistance profiles of the adults and other traits, including longevity, egg laying and pupation. Generational larval exposure (*An. gambiae*) to heavy metals such as cadmium (Muturi et al., 2018) was found to induce the overexpression of several genes involved in detoxification, transport and protein synthesis. Similarly, *An. coluzzii* larvae collected from agricultural breeding sites contaminated with copper demonstrated high resistance to λ-cyhalothrin with a positive correlation between the resistance levels and copper pollution levels (Talom et al., 2020).

Exposure to environmental pollutants may activate similar chemo-protective mechanisms as insecticides thus augmenting selection pressures leading to metabolic resistance in disease vectors (Sadia et al., 2024), although this is poorly researched (Nkya et al., 2013). Polycyclic aromatic hydrocarbons (PAHs) are one of the most recalcitrant and ubiquitous environmental pollutants, derived from various activities, including the incomplete combustion of organic matter (Abdel-Shafy and Mansour, 2016). PAHs are known to act as potent ligands of the aryl hydrocarbon receptor (Ahr) in higher organisms, whose activation leads to the overexpression of various detoxification enzymes, including the cytochrome P450s, glutathione S-transferases (GSTs), and carboxylesterases (Bohonowych and Denison, 2007; Soshilov and Denison, 2014). PAHs are metabolised in cells via the Ahr-dependent or Ahr-independent pathways. The former involves core detoxification genes of the cytochrome P450, GST, and epoxide hydrolases families, whereas the latter involves the use of aldo-keto reductases that often lead to the activation of PAHs to more reactive intermediates (Goedtke et al., 2021; O’Driscoll et al., 2018; PHE Centre for Radiation, 2008). The involvement of the Ahr in the transcriptional regulation of P450 monooxygenases through promoter and enhancer sequences in insects has been well documented (Pan et al., 2019; Peng et al., 2017; Vogel et al., 2020).

The impact of PAH pollutants on mosquito Ahr-activated metabolic genes is poorly understood. Some studies have investigated genes activated by exposure to PAHs in insects (David et al., 2010; Zhang et al., 2019). Mosquitoes that survive in PAH polluted breeding sites have been shown to have elevated levels of detoxification enzymes, including the cytochrome P450 monooxygenases, GSTs, uridine-diphospho-glucuronosyl transferases (UGTs) and esterases (ref). leading to significant physiological changes (Tetreau et al., 2014). Such genes could potentially be involved in metabolic cross-resistance towards insecticides. Unfortunately, little is known about the detoxification enzymes involved in cross-resistance between the PAHs and insecticides in mosquitoes. To bridge this gap, we conducted a series of multi-generational lab selections with PAHs, fluorene, naphthalene and a mix of both using field and laboratory-susceptible strains of *An. coluzzii,* followed by insecticide-resistance phenotyping, whole transcriptome analysis using RNASeq and *in vitro* functional characterization to understand the impact of PAHs in the evolution of insecticide resistance. Several genes and gene families earlier implicated in insecticide resistance were found to be associated with exposure to PAHs. The P450 *CYP6M4* was the most consistently overexpressed metabolic gene in all the selected lines. It was expressed in *E. coli* and shown to metabolize pyrethroids and PAHs, thereby confirming the cross-resistance potential of selection by environmental pollutants on resistance towards public health insecticides.

## 2.1 Materials and Methods

### 2.1.1 Mosquito populations

Blood-fed female mosquitoes, resting indoor were collected, between August and October 2018, using handheld battery-powered aspirators (John. W. Hock, Florida, USA) in Auyo (12°21′N, 9°59′E), a town in northern, Nigeria, with extensive irrigation activities (Ibrahim et al., 2014). Mosquitoes were morphologically identified (Gilles and Coetzee, 1987) as *An. gambiae sensu lato* and forced to lay eggs in 1.5 ml Eppendorf tube (Morgan et al., 2010). The F_0_ parents and egg batches were transferred to Liverpool School of Tropical Medicine under a DEFRA Licence (PATH/125/2012). Genomic DNA was extracted (Livak, 1984) and used to identify the F_0_ parents to species level using SINE_200 PCR (Santolamazza et al., 2008), before eggs were pooled in trays and maintained under standard insectary conditions (25°C, 75% relative humidity and 12h:12h light:dark cycles). Subset of the F_1_ females that emerged were mixed randomly, and 3-5 d females were used for insecticide susceptibility assays. Egg batches from females of the laboratory susceptible colony, Ngousso (Harris et al., 2010) were also obtained from the Liverpool Insect Testing Establishment (LITE) for the subsequent experiments described below, which were conducted in parallel with the field population mentioned above.

### 2.1.2 Preliminary determination of diagnostic LC_50s_ for PAHs assays

Lower members of the PAHs with fewer ring structures and no substitution, which make them more persistent in the environment (Baldwin et al., 2020) and not easily undergoing microbial metabolism (Kanaly and Harayama, 2000) were chosen as representative PAH for the selection experiments. These included fluorene and naphthalene, which were purchased from SIGMA (Sigma Aldrich, Schnelldorf, Germany). These PAHs were dissolved separately in absolute ethanol to a final concentration of 1g/100 ml (stock concentration). A series of concentrations (in 100 ml of distilled water), 0, 3, 6, 12, 15, 20, and 25 μg/ml, were prepared from the above stock solution. Five replicates of fourth instar larvae (25 each) and a control group (n = 25) were exposed to PAHs in paper cups containing 100 ml of distilled water for 24 h (WHO, 2005). Mortality was calculated in each of the treated groups and compared with the control group exposed to only water containing the same volume of ethanol. Dose-response curves were created using probit analysis (Finney, 1964) from which the LC_50_ (concentration that kills 50% of the population) was calculated for each PAH. The calculated LC_50_s for each fluorene and naphthalene were used for the selection experiment in which the same exposure procedure described is repeated, only with more replicates, such that at each generation, enough survivors that maintained the line were obtained from the various exposures.

### 2.1.3 Determination of the susceptibility of adults to insecticides using WHO conventional and synergist bioassays

Bioassays were conducted according to the WHO protocol (WHO, 2016) to determine susceptibility to insecticides. Briefly, four replicates consisting of 25 females (all 3-5 d old) were exposed to permethrin (0.75%), deltamethrin (0.05%), DDT (4%), bendiocarb (0.1%) and malathion (5%) insecticides for 1 h. One replicate of 25 mosquitoes from the same colony and age was used as a control exposed only to papers impregnated with the carrier oil, respectively. The mosquitoes were transferred to holding tubes for recovery, where they were fed with a 10% sucrose solution for 24 h, after which mortalities were recorded (WHO, 2016).

To investigate the role of cytochrome P450s in resistance of the field *An. coluzzii* at the tenth generation, synergist bioassay was conducted using piperonyl butoxide (Feyereisen, 2015). Four replicates each of 25 females were exposed to 4% PBO impregnated papers for 1 h before transferring them to tubes lined with 0.75% permethrin impregnated papers for another 1 h. Mosquitoes were immediately transferred to holding tubes for 24 h after which the mortality was recorded. Two separate tubes, each with 25 females were used as control, the first one containing only 4% PBO, while the second tube contained 0.75% permethrin.

All the insecticide, synergist, and control papers were obtained from the WHO certified source (Vector Control Research Unit, Universiti Sains Malaysia).

### 2.1.4 Multi-generational selection of field (Auyo) and laboratory (Ngousso) populations using PAHs

A second set of Auyo egg batches described above was raised to the F_1_ progenies and used for selection experiments. The F_1_ individuals were divided into 4 different lines: (i) line 1, control (non-selected control, NSC), which were maintained in the insectary for 10 generations without any selection; (ii) line 2, selected with fluorene (AnFluo); (iii) line 3, selected with naphthalene (AnNaph); and (iv) line 4, selected with 1:1 mixture of fluorene and naphthalene (AnMix), which allowed to investigate the impact of simultaneous exposure to the above two PAHs. The laboratory susceptible *An. coluzzii* colony, Ngousso, obtained from Liverpool Insect Testing Establishment (LITE) was also divided into three lines. Line 1 was the control, maintained in the insectary without selection (Control). Line 2 and 3 were selected on naphthalene and fluorene, respectively (Ngou-Naph and Ngou-Fluo). More details of the mosquito lines are provided in Table S1. Fourth instar larvae from each of these lines (25 per replicate) were exposed to the respective PAHs in paper cups for 24 h, while the control groups were exposed only distilled water containing the same volume of ethanol used to dissolve the PAHs. Mortalities were recorded, and larvae alive after 24 h were washed in distilled water and transferred to new trays and fed with tetramin® baby fish food until adult emergence. The emerged adults were blood-fed after 3-4 d after and allowed to lay eggs, and form the subsequent generations which were used subsequently for selection with the higher concentrations of PAHs. Details of the concentrations used at each generation are provided in Table S2, additional file 2. Insecticide susceptibility tests were conducted using the WHO tubes protocol (WHO, 2016) with 0.75% permethrin, 0.05% deltamethrin, 0.1% bendiocarb, 4% DDT and 5% malathion using selected females at the F_0_, F_7_ and F_10_ generations. For the laboratory susceptible colony selections, selected individuals at F_10_, unexposed to insecticides and from the non-selected control line, also unexposed to insecticides, were sampled and kept in freezers for RNA extraction/transcriptional analysis. For the field selection lines, survivors of bioassay (permethrin 0.75%) at F_10_ and those unexposed to insecticides from selected and non-selected controls at the tenth generation required for transcriptional analysis were kept in the freezer (−80 °C) until needed for RNA extraction.

### 2.2 RNAseq Genome-wide Transcriptomics Analysis

#### 2.2.1 RNA extraction and library preparation and sequencing

RNA was extracted from 3 replicates of pools of 10 permethrin selected female mosquitoes (surviving exposure at the 10^th^ generation) and unexposed lines (Table S1, additional file 2). Total RNA was extracted using the Arcturus PicoPure kits (Applied Biosystems, CA, USA) according to the manufacturer’s instructions and the RNA was treated with DNase (Qiagen, Hilden, Germany) to remove the residual genomic DNA. The concentrations and integrity of the isolated RNA were determined using Agilent 4200 TapeStation (Agilent Technologies, CA, USA).

Quality control, cDNA library preparation and sequencing of the RNA were conducted at Greta Immobile Molaro, Polo d’Innovazione di Genomica Genetica e Biologia SCaRL, Italy, under the sponsorship of Research Infrastructure for the control of vector-borne diseases: Infravec2 supported by Horizon 2020 (Grant Agreement No. 731060). Illumina chemistry V2.5, 2x75 bp paired end running was employed in the RNA sequencing using the NextSeq 500 machine. Libraries were prepared in the same laboratory using TruSeq Illumina mRNA Stranded Sample Preparation for Illumina Paired-End Indexed Sequencing. Qubit 4.0 Fluorometer (Thermo Scientific, MA, USA) was used to determine the concentrations of the libraries.

#### 2.2.2 RNAseq data analysis

Analysis was conducted using a Snakemake-based pipeline (*RNA-Seq-Pop*) (Ibrahim et al., 2023; Nagi et al., 2023), a suite which enables differential gene expression analysis, identification of single-nucleotide polymorphisms, and population genetics analysis. Briefly, fastq files of all the paired results, reference transcriptome of *An. gambiae* AgamP4.10 (downloaded from Vectorbase) and sample metadata are provided for the workflow. Raw fastq reads were trimmed after the quality control is conducted using Cutadapt (v4.0) (Martin, 2011). Reads with low-quality and minimum score <20 were removed. This is followed by the alignment of the trimmed reads with the reference transcriptome using KALLISTO (Bray et al., 2016), and differential expression at the gene level conducted using DESeq2 (Love et al., 2014), and at the isoform level using Sleuth (Pimentel et al., 2017). Differential expression analysis was conducted for between the selected and the non-selected control of both the field and laboratory strains. Comparisons were also made between the selected and non-selected field lines with the baseline control (Ngousso control) and between exposed and unexposed to permethrin groups within the Auyo (field) strain. Log_2_ fold change >1 was imposed for all these comparisons while maintaining a 0.05 false discovery rate (FDR) level of significance. For data visualization, principal component analyses (PCA) were created using differentially expressed genes between replicates and treatments. Volcano plots were also created using the most differentially expressed genes, as well as heatmaps for the most overexpressed genes (Blighe et al., 2018).

#### 2.2.3 Validation of overexpressed candidate genes using qRT-PCR

To validate the expression levels of select candidate genes found to be overexpressed in the RNASeq and previously associated with insecticide resistance, quantitative real-time PCR was conducted using RNA samples from RNAseq experiments above. cDNA was synthesized from 1 μg of total RNA as a template, using oligo (dT) primers and superscript III transcriptase enzymes (Invitrogen, Waltham, CA USA) according to the manufacturer’s instructions. Primers utilised are provided in Table S3 (Additional file 2). These include cytochrome P450s: *CYP6Z1, CYP6Z3, CYP6M4, CYP6M3, CYP6Z2* and *CYP6P3*; GSTs: *GSTe2, GSTe5,* and *GSTe4;* aryl hydrocarbon receptor (Ahr) and its nuclear counterpart aryl-hydrocarbon receptor nuclear translocator (*AHRNT/Tango);* the aryl hydrocarbon receptor-interacting protein (AIP); and the co-chaperones heat shock protein 90 (*HSP90).* Elongation factor α (ELF) and ribosomal protein S7 (RSP7) were used as housekeeping genes for normalization of expression. The reaction proceeded with 1 μl of the cDNA as template in which 10 μl of SyBr Green fluorescent dye (Sigma Aldrich, Germany), 0.6 μl each of forward and reverse primers were added. The reaction was topped up to 20 μl with the addition of 7.8 μl nuclease-free water. The thermocycling conditions consisted of 95 °C (initial denaturation) for 3 min, followed by 40 cycles each at 95 °C for 10 s and 60 °C for 10 s (Muhammad et al., 2021).

The Cq values generated from the run were used to calculate the normalised expression levels of the mRNA in each sample by comparing the values in our genes and the housekeeping genes. The efficiencies of the primers were also incorporated into the calculations, after which values were converted to their logarithmic forms for normal distribution according to the established protocol of ddCT (Schmittgen and Livak, 2008).

#### 2.2.4 Investigation of the signatures of selection due to generational PAH exposures

Using the *RNA-Seq-Pop,* SNP variants were called using the generated alignment files (BAM format) for each treatment as a separate population with the Bayesian haplotype-based caller *freebayes* version 1.3.2 at the ploidy level of 20 (10 individual mosquito pools) (Garrison and Marth, 2012). The called variants were later annotated using *snpeff* version 5.0 (Cingolani et al., 2012). Through applying a missingness proportion of 1 and quality score of 30, the called variants were used to perform a windowed Hudson’s F_ST_ scan for each pairwise comparison (Bhatia et al., 2013; Hudson et al., 1992).

### 2.3 Functional characterization of *CYP6M4*

#### 2.3.1 Preparation of recombinant CYP6M4 for *in vitro* expression

Primers were designed to amplify the full length of *CYP6M4* (Table S4, Additional file 2) using Phusion Hot Start II Taq polymerase (Thermo Scientific, MA, USA) and 3 replicates of the cDNA from Auyo population as templates. The amplified fragments were purified using Qiaquick purification kits (Qiagen, Hilden, Germany) and cloned into pJET1.2 blunt cloning vector, transformed into *DH5α* cells (Thermo Fisher Scientific, MA, USA) and mini-prepped. Fifteen (15) clones were sequenced to establish the dominant allele of the *CYP6M4*. Full-length coding sequence of the dominant allele was fused to the amino terminal of bacterial *ompA+2* leader sequence with its associated downstream ala+pro linker using Phusion Hot Start II Taq Polymerase PCR according to the previously established protocols (Pritchard et al., 1997; Riveron et al., 2013). These were digested out from pJET1.2 vector using *Xba*I and *Nde*I enzymes, purified using Qiagen gel extraction kit ((QIAGEN, Hilden, Germany) and ligated into pCWori+ linearised with the same enzymes. The pCWori+CYP6M4 construct was co-transformed together with pACYC-184 plasmid bearing *An. gambiae* cytochrome P450 reductase (CPR) into the *E. coli* JM109 competent cells (All primers provided in Table S4, additional file 2) for subsequent co-expression as described earlier (Ibrahim et al., 2015; Pritchard et al., 1997)

#### 2.3.2 Heterologous expression of recombinant CYP6M4

The CYP6M4 recombinant protein was expressed and purified following the procedures described earlier (Ibrahim et al., 2016; Pritchard et al., 1997). A starter culture supplemented with chloramphenicol and ampicillin (to the final concentrations of 34 μg/ml and 50 μg/ml, respectively) was inoculated with JM109 cells transformed with the above P450 and reductase plasmids and left to shake for 12-14 h. 200 ml of pre-heated terrific broth (to 37 °C) (Alpha Teknova, Hollister, USA) was inoculated the following day with 2 ml of the starter culture and allowed to shake at 200 rpm and 37 °C until absorbance of ∼0.6-0.8 was reached. Expression was induced by the addition of 1 mM isopropyl β-D-1-thiogalactopyranoside (IPTG) and 0.5 mM *δ-*aminolevulinic (ALA) to the final concentrations. For optimum expression conditions, shaking at 150 rpm and 25°C was applied and left overnight, while monitoring started at ∼13 h after induction.

#### 2.3.3 *In vitro* metabolism assay with insecticides and PAHs

Metabolism assays were carried out using 10 μM naphthalene, fluorene and fluoranthene, as representative PAHs, and insecticides from three different classes, including permethrin, α-cypermethrin, and deltamethrin, bendiocarb, propoxur, and DDT (10 μM of each). Assay protocols were as described previously (Ibrahim et al., 2024, 2018). The recombinant protein (50 picomol) comprised of CYP6M4 protein and *An. gambiae* CPR was constituted into 0.1M potassium phosphate buffer in a 1:10 ratio with the *cytochrome b5* protein. The composition of the enzyme-buffer mix and that of the regeneration mixture were as described earlier (Ibrahim et al., 2018). Reactions were initiated through the addition of the NADPH regeneration mixture (containing NADP^+^) and insecticides. As controls, similar reactions were carried out but with the regeneration buffer devoid of NADP^+^ (-NADPH). The mixes were allowed to shake for 2 h at 1200 rpm and 30°C. To stop the reactions, ice-cold acetonitrile (100 μl) was added to all the tubes and the tubes were shaken for 5 min before centrifuging at 13,000 rpm for 15 min. The supernatants (150µl) were loaded into HPLC vials for injection. For the pyrethroids and DDT, a mobile phase consisting of 70:30 acetonitrile and water, respectively and run at a flow rate of 1 ml/min were used, with detection wavelength set to 226 nm. Bendiocarb and propoxur were detected at 232 nm wavelength with a mobile composition of 60:40 acetonitrile and 0.1% phosphate acid in water. Polycyclic aromatic hydrocarbons were detected with 80:20 acetonitrile: water mobile phase, wavelength of 254 nm, and a run for 20 min,with a flow rate of 1ml/min (Onopiuk et al., 2022).

#### 2.3.4 Determination of steady-state kinetics parameters

To study the reaction kinetics of the metabolism of α-cypermethrin and permethrin by the CYP6M4, assays were conducted again with 50 pmol of recombinant membrane protein, and available substrate concentrations (2.5 to 25 μΜ) for 30 min incubation period. The reactions were carried out in triplicates, and the resulting data were used in the calculation of kinetic parameters (K_m_ and V_max_) using the plots of substrate concentrations against the initial velocities. Using the least squares non-linear regression, the data were fitted to the Michaelis-Menten module plots, allowing for the calculation of the kinetic parameters. The plots were created using GraphPad Prism version 8.0.1 (GraphPad Inc., La Jolla, CA, USA).

#### 2.3.5 Sequence characterization of *An. coluzzii* CYP6M4 and prediction of structurally conserved regions

To predict the conserved features of *CYP6M4*, which could impact its activity, its coding sequence was compared to other closely related P450s. These P450s include *An. gambiae CYP6M2* (AGAP008212), which shares 98.2% identity with *An. coluzzii CYP6M4* (Nikou et al., 2003) and another ortholog *An. funestus CYP6M4* (Afun019401), which shares 79.6% identity with *An. coluzzii CYP6M4* (Irving et al., 2012). Putative substrate recognition sites 1-6 of *CYP6M4* and the above P450s were compared by mapping their amino acid sequences to those of *Pseudomonas putida CYP101A* (P450cam) (Poulos et al., 1985) and structurally conserved regions of the P450s were predicted using an online tool, CYPED (Sirim et al., 2010). Figure S6 was prepared using the CLC sequence viewer 7.0 (http://www.clcbio.com/).

#### 2.3.6 Prediction of the role of *An. coluzzii CYP6M4* in pyrethroid and PAHs metabolism using homology modelling and docking simulations

To predict the folding pattern of CYP6M4 and identify the role of its active site amino acid residues in metabolism of pyrethroids and PAHs, a homology model of the predominant sequence used for functional validation was created using the Modeller 10.5 (Eswar et al., 2005) with the crystal structure of human *CYP3A4* [PDB: 1TQN, (Yano et al., 2004) which share 31.7% identity with *An. coluzzii* CYP6M4, as a template. A total of 20 models were generated for each sequence, and the models were assessed externally using Errat version 2 (Colovos and Yeates, 1993) to identify the best models from statistical patterns of non-bonded interaction between different atom types. Virtual datasets of ligand insecticides: 1R-cis permethrin (ZINC01850374) and α-cypermethrin (ZINC71789490) were retrieved from ZINC12 database (Irwin and Shoichet, 2005); while fluorene (CID_6853) and fluoranthene (CID-9154) structures were retrieved from ChemID plus (https://pubchem.ncbi.nlm.nih.gov/source/ChemIDplus). Docking simulations were carried out using the Achilles Blind Docking Server (Sánchez-Linares et al., 2012), which uses Vina_vision – a customised version of Autodock Vina. For each ligand, 30 binding poses/clusters were generated and sorted according to binding energy and conformation in the model’s active site. Figures were prepared using the PyMOL 2.4 (DeLano and Bromberg, 2004). Non-bonded interactions were predicted using the protein-ligand interaction profiler (Salentin et al., 2015).

## 3.0 RESULTS

### 3.1 Resistance profiles of mosquito colonies and their tolerance to select PAHs

The field, F_1_ *An. coluzzii* was highly resistant to deltamethrin, permethrin and DDT, with mortalities of 11%±1.7, 16.3%±2.1 and 13.8%±2.9, respectively (Figure 1a). Moderate resistance was observed towards bendiocarb (mortality = 93%±0.4) and there was a full susceptibility towards malathion (99%±0.3). The larvae of the field population exhibited higher naphthalene tolerance (LC_50_ of 8.8μg/l ±1.3), compared with fluorene, with LC_50_ of 4.9μg/l ±10.1, with the highest susceptibility observred in the larvae exposed to fluoranthene (LC_50_ of 1.8μg/l ±0.1) (Fig. 1b).

**Figure 1:**
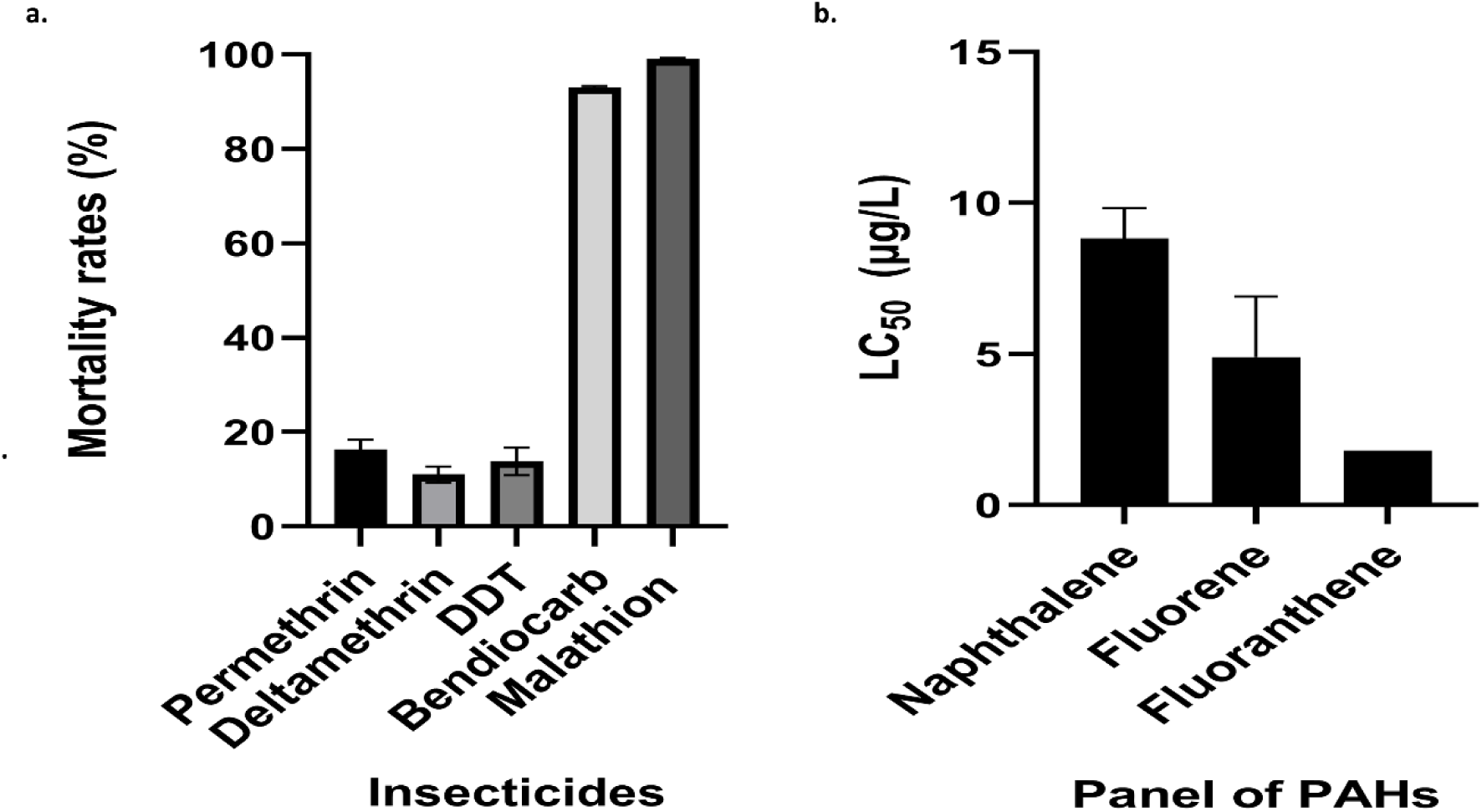
Pre-selection insecticide and PAH susceptibility profiles of the Auyo *An. coluzzii*. (a) Mortality rates of the adult mosquitoes exposed to diagnostic doses of insecticides. Values are means ± standard deviations of 4 replicates each of 25 females (b) The LC_50_ of the select PAHs on the larvae of the field population used in the selection experiments. Values represent mean ± standard deviations of 10 replicates of 30 larvae each

#### 3.1.1 PAH selection of field resistant *An. coluzzii* induced progressive increase in insecticide susceptibility

To investigate the potential role of PAHs on resistance the field resistant *An. coluzzii*, resistant towards insecticides, was subjected to selection with PAHs, naphthalene, fluorene and the mixture of both (AnNaph, AnFluo and AnMix) for ten generations. Resistance profiles were monitored using WHO tube bioassays at the seventh and tenth generations. The field mosquitoes control line was maintained in the insectary with no selection. A progressive increase in susceptibility for both selected and non-selected lines were observed, particularly for permethrin, deltamethrin and DDT. While mortality following exposure to the diagnostic dose of permethrin at F_1_ (Fig. 2a) and deltamethrin (Fig. 2b) were 17.0%±3.3 and 11±3.2%, respectively, these mortalities progressively increased to 78%±4.4 (*P* = 0.001) and 100% (*P* = 0.0001) at the tenth generation, respectively (Fig. 2 a and b). Similar trend was observed in the case of DDT with F_1_ mortality of 19%±4.4, that rose to 81%±15.6 (*P* = 0.0003) at the tenth generation for the naphthalene selected line (Fig. 2c). No significant differences were observed when comparing the tenth-generation mortality rates of permethrin of the selected AnNaph (78%±4.4, *P* = 1.000), AnFluo (73%±11.4, *P*= 0.4671)) and AnMix ((82%±4.5, *P* = 0.3272) with non-selected control lines (NSC). However, selection with fluorene showed a significantly lower deltamethrin mortality at the seventh generation (*P= 0.051)*, indicating a potential role of fluorene in increasing resistance to deltamethrin. In contrast, at the tenth generation, selection with naphthalene led to significantly higher deltamethrin mortality than in the non-selected control lines (AnNaph mortality = 100%, *P* = 0.001, NSC mortality = 88%±2.0). These findings suggest that longer (up to 10 generations) exposure of this population to PAHs increased deltamethrin susceptibility at the tenth generation, but with no significant impact on permethrin’s susceptibility at the tenth generation. However, higher permethrin mortality was recorded for the non-selected control compared to the fluorene selected line (AnFluo) at both the seventh and tenth generations (not statistically significant). In the case of DDT, at the seventh generation, the selected line demonstrated significantly (*P* = 0.01) lower mortality rates compared to the non-selected control, suggesting the potential of PAH exposure to select for DDT resistance. In contrast, at the tenth generation, the naphthalene selected line (AnNaph) showed significantly (*P = 0.001*) higher mortality compared to the non-selected line (Fig. 2c), whereas the fluorene selected line (AnFluo) showed significantly lower mortality than in the control (*P= 0.0023*), indicating how the PAHs can potentially have different/contrasting impacts on resistance. Bendiocarb mortality increased significantly *(P = 0.*002) from 93%±1.7 for F_1_ to 100% for the seventh and tenth generations (Fig. 2d). and there were no changes in mortalities for malathion observed (Fig. 2e).

**Figure 2:**
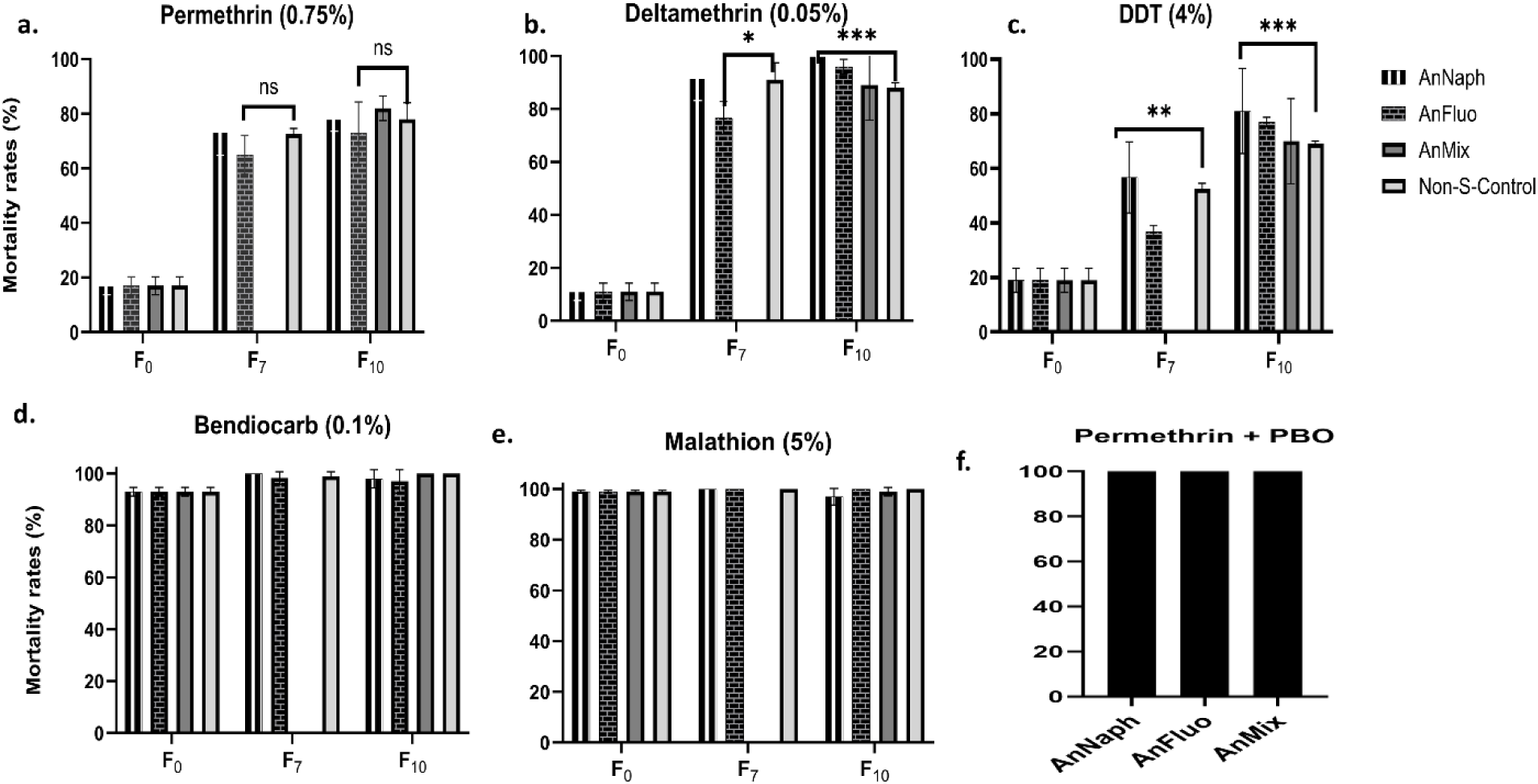
Susceptibility profiles of the various PAH-preselected experimental lines of *An. coluzzii* field population (Auyo) following exposure towards public health insecticides. Bioassays conducted using WHO diagnostic doses of the different insecticides conducted on the adult female (a., b., c., d. and e). Synergist assay conducted using PBO to investigate the role of cytochrome P450s in the recovery of susceptibility to permethrin in the selected lines (panel f). Results are presented as the mean of the mortality ± standard deviations of the means.

To determine the potential contribution of P450-based metabolic resistance towards the residual resistance, colonies were exposed to insecticides in the presence of PBO. PBO fully restored susceptibility to permethrin (100% mortality) (Fig. 2f).

The laboratory susceptible strain Ngousso was also subjected to selection with fluorene and naphthalene for 10 generations, with bioassays conducted after the 7^th^ and 10^th^ generations to monitor changes in resistance in comparison to the non-selected control line of the same strain (Ngousso). Permethrin mortality for the naphthalene-selected line (Fig. 3a) was significantly (*P= 0.039*) lower than the mortality recorded for the non-selected line, indicating strongly that naphthalene exposure for seven generations led to the development of resistance to permethrin. However, the permethrin mortality at the seventh (*P= 0.2682*) and tenth (*P=* 0.1156) generations for the fluorene selected line were slightly lower than the mortality recorded for the non-selected control lines. Furthermore, the mortality rates for DDT (Fig. 3c) in the selected lines were significantly lower than those of the non-selected lines at the 7^th^ and 10^th^ generations (*P* = 0.0128), indicating the potential of PAHs in causing cross-resistance to DDT in a controlled experiment. Furthermore, slower knockdown rates were observed in the selected lines compared to the non-selected control for all the insecticides tested except deltamethrin, which had slightly faster rates in the selected lines. Contrary to these, the mortality rates for deltamethrin, bendiocarb and malathion (100%) remained the same throughout the selection regime with no observed changes in the selected and non-selected control lines (Fig. 3 b, d, and e).

**Figure 3:**
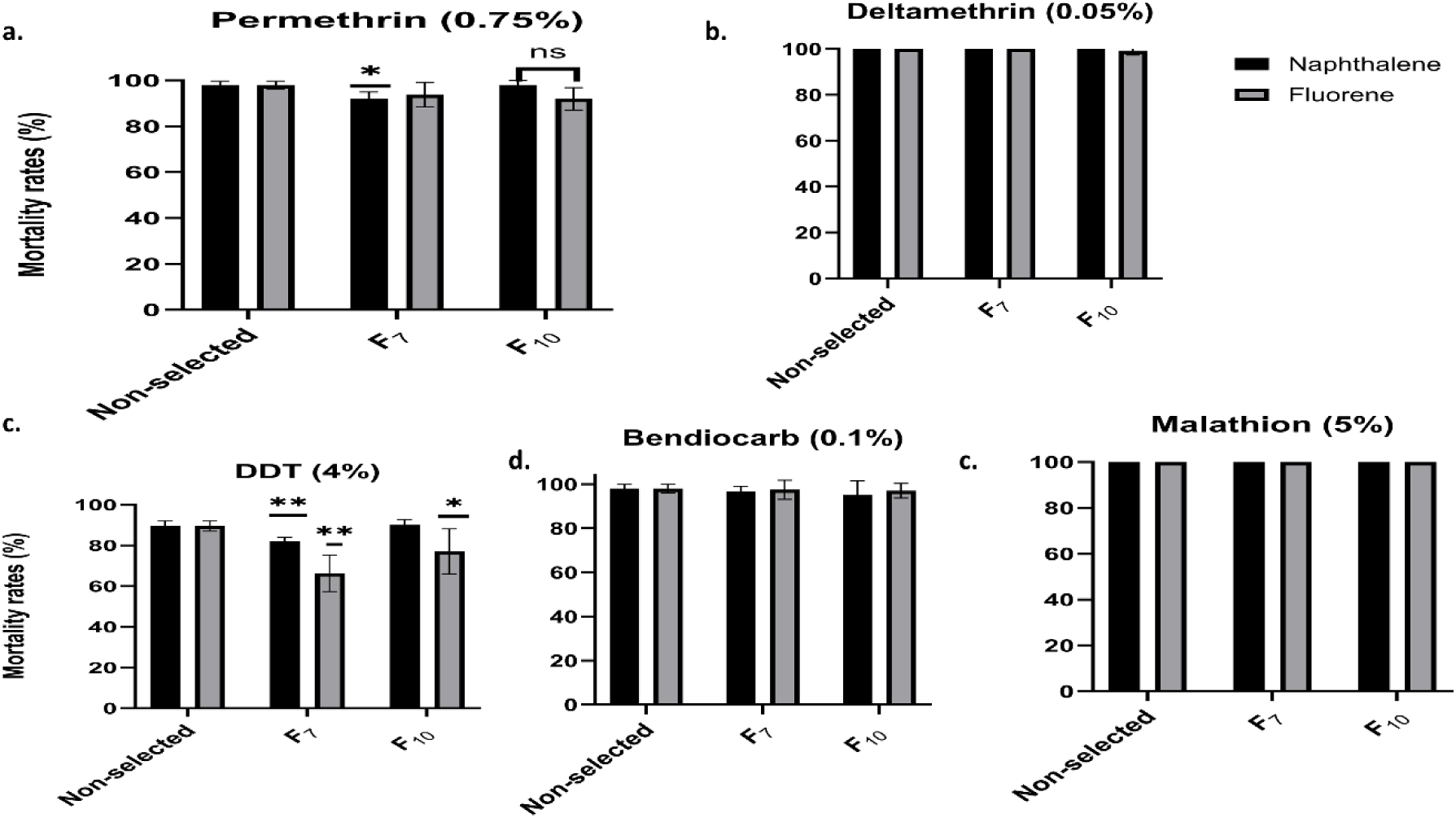
Susceptibility profiles of the various PAH-preselected experimental lines of *An. coluzzii* laboratory susceptible colony (Ngousso) following exposure towards public health insecticides. Bioassays were conducted with the WHO diagnostic concentrations for each insecticide. Values are presented as mortality mean ±standard deviations of the means

### 3.2 PAH selection modifies the expression of metabolic genes in the field (Auyo) and laboratory susceptible (Ngousso) strains of *An. coluzzii*

#### 3.2.1 Differential expression profiles of genes associated with fluorene selection

##### 3.2.1.1 Auyo field populations

Selections with fluorene led to the overexpression of several gene families earlier implicated in insecticide resistance and various other physiological processes in *Anopheles* species (Table 1). The top commonly overexpressed in the comparisons, R vs NSC and C vs NSC (R refers to survivors of permethrin exposure at the tenth generation while C is the unexposed to permethrin at the 10^th^ generation, considered as control, NSC stands for the non-selected control line kept for ten generation with no selection on PAHs used as the background control) genes include (with high fold changes in the exposed vs NSC) An immune gene (AGAP029286) (FC = 451), Protein spinster (AGAP006345) (FC= 139), CYP6S2 (AGAP008203) (FC =17), Chymotrapsiongen B (AGAP029120) (FC= 13), Extracellular protease inhibitor 10 (AGAP029163) (FC = 11.3) and Trypsin (AGAP001251) (FC = 9). Other highly expressed genes include fatty acyl-CoA reductase1, several odorant-binding proteins (including 47, 49, 83a and its receptor), immune and sodium-independent sulfate anion transporter. Several other genes directly involved in insecticide resistance due to their detoxification nature were also found to be commonly overexpressed following generational exposure of the field population to fluorene. Notable among these include *CYP6M4 (FC = 5.1)* and *CYP4C7 (FC = 4.5)* (Table 1). The highly overexpressed and downregulated genes in the various comparisons have been plotted in volcano plots with some of the genes of interest (e.g. *CYP6M4*, distinct red dot) shown (Fig. 4). Other major detoxification genes overexpressed in at least one comparison are depicted in heatmaps (Figure S3, additional file 1).

**Figure 4:**
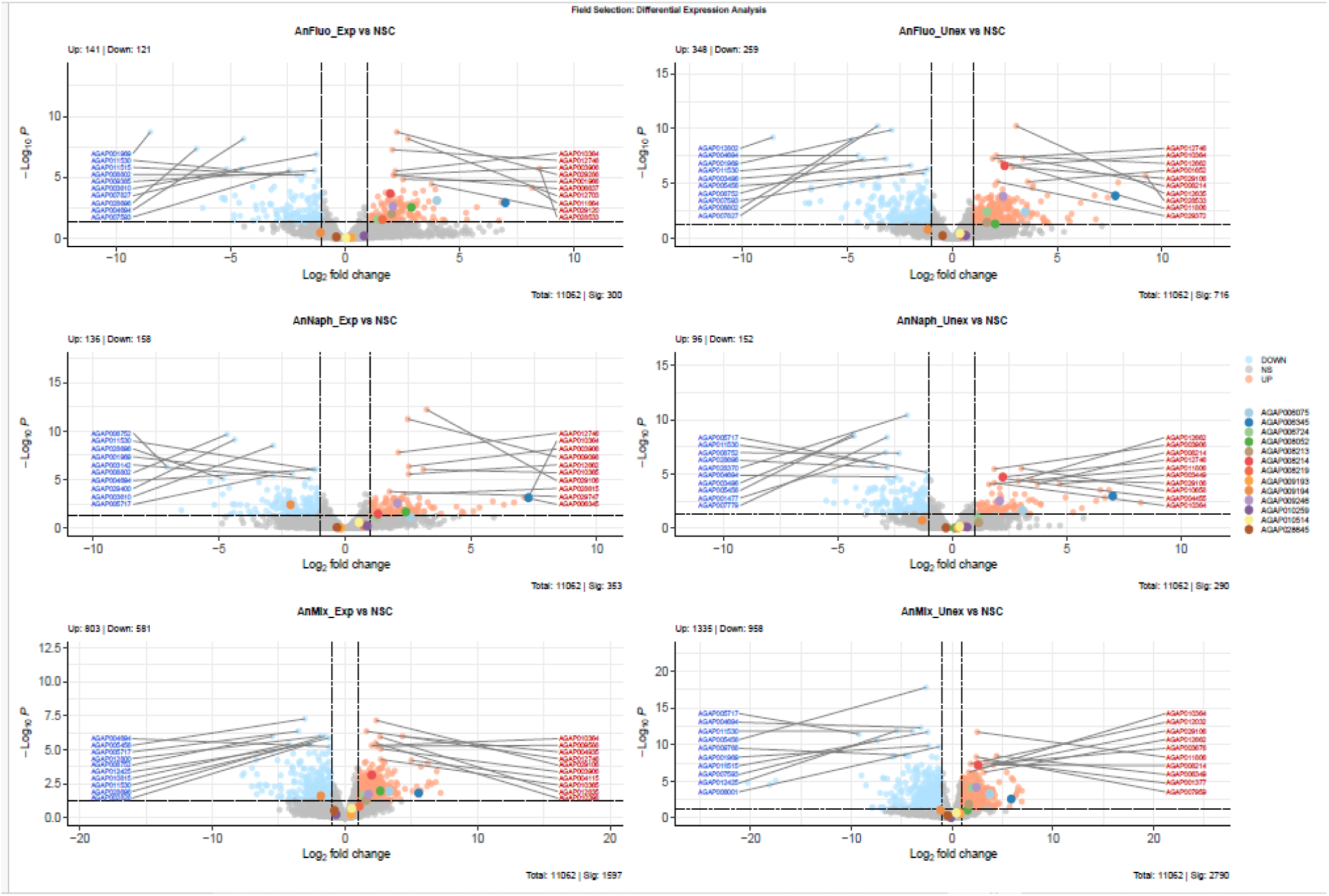
Differentially expressed genes of interest in the different selected lines. Volcano plots top genes of interest differentially expressed, showing the log2 fold change (x-axis) and significance level (−log10 P value) on the y-axis, highlighting the significantly up and down-regulated genes. Some genes of interest are also highlighted with the CYP6M4, notably shown as red dot throughout the plots to indicate consistent overexpression in different selection lines. Comparison was made between the selected lines (exposed and unexposed to permethrin diagnostic doses in a contact bioassay) and the non-selected control (NSC) line.

**Table 1:**
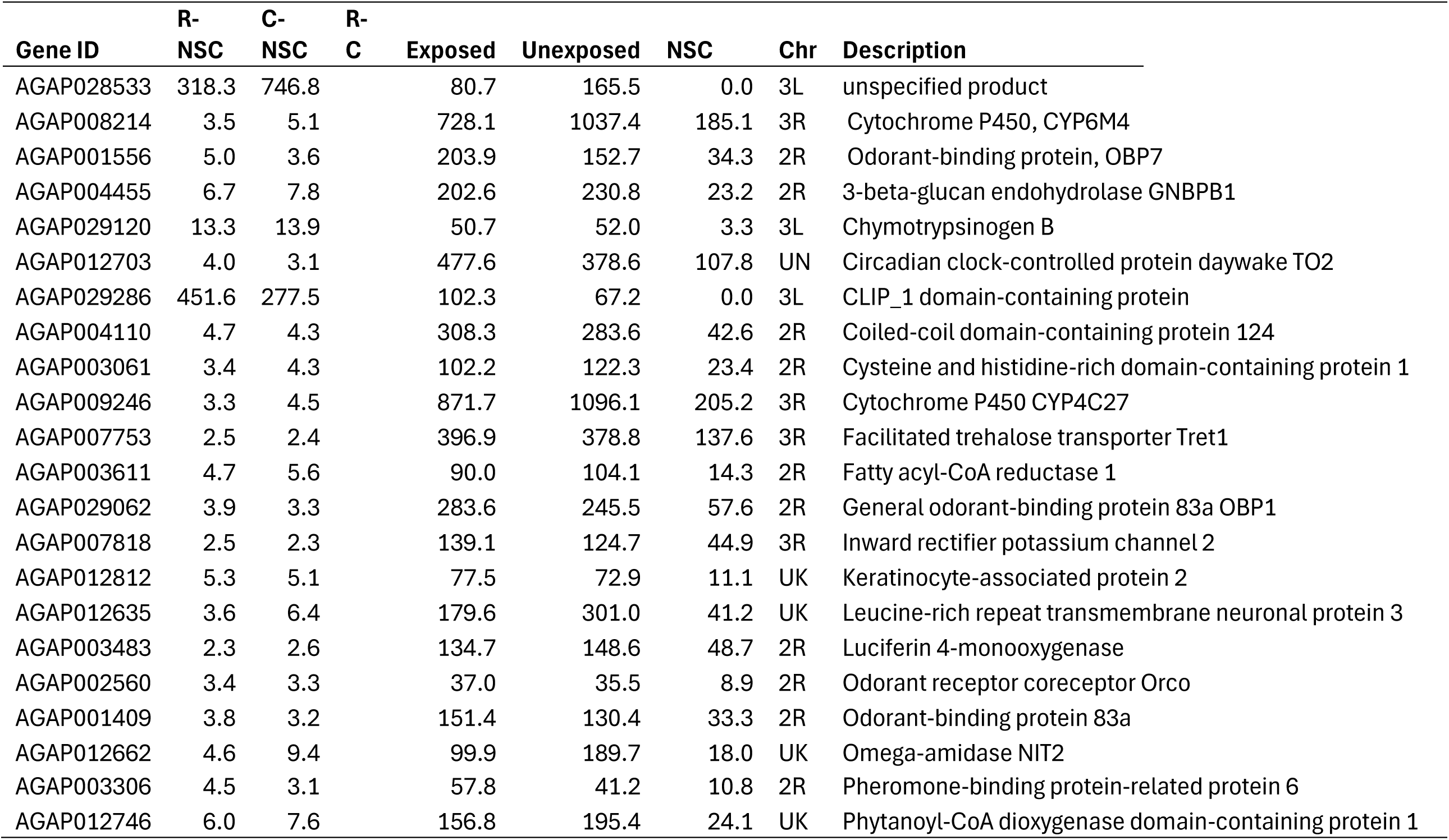
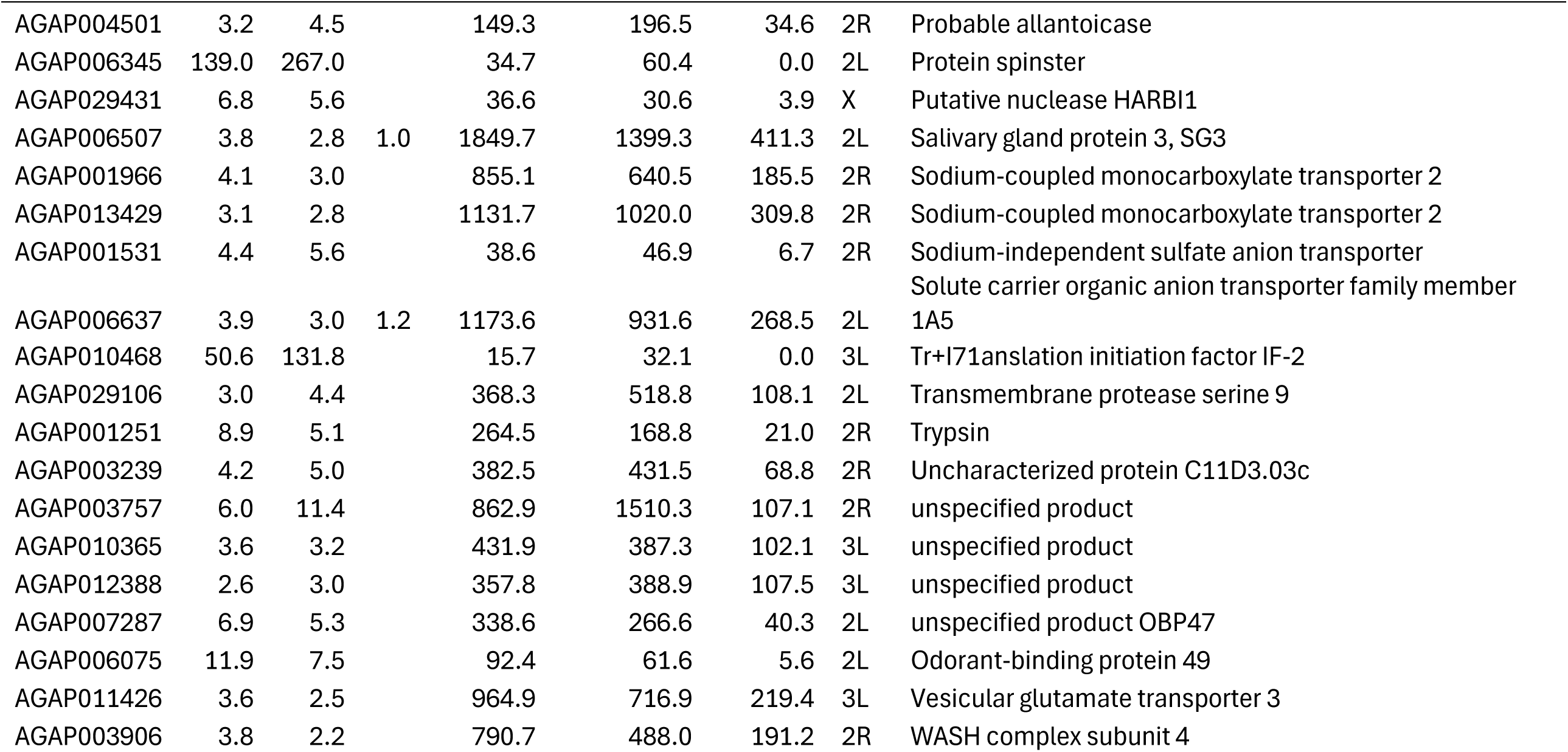
Commonly overexpressed genes in the field population selected on fluorene (AnFluo) compared to field non-selected control (NS) line. Exposed (R) survived permethrin bioassays at the tenth generation of selection, while the unexposed (C) were from the same selected lines but no insecticide exposure**. (log_2_FC values >1 and FDR-adjusted *p* < 0.05)**

##### 3.2.1.2 Ngousso Laboratory susceptible strain

The highly overexpressed genes in the Ngousso fluorene (Table S7) selection include: The 70kDa heat shock protein (AGAP004581, FC = 74), *CYP6N1* (AGAP008210, FC= 10), ZP domain-containing protein (AGAP000786, FC = 10), leucine-rich immune protein (Coil-less) (AGAP002542, FC = 9.5), cuticular protein RR-1 family 20 (*CPR20*, AGAP005969, FC = 7.7), Ser/Thr protein phosphatase/nucleotidase (AGAP005458, FC = 8.9), chitinase (AGAP011033, FC = 4.7), Alkylated DNA repair protein alkB homolog 8 (FC = 6.7), Plasma kallikrein (FC=6.1), and Ecdysteroid UDP-glucosyltransferase (AGAP009137, FC = 4.5).

For the selection of Ngousso with both fluorene and naphthalene there a was a common overexpression of genes relating to, i) behavioural proteins (including Gustatory receptor and protein takeout, ii) cuticular proteins from all classes (e.g *CPR20*, CPR75, and *CPLCP25*), iii) digestive enzymes (like the proteases, chitinases, chymotrypsinogen and so on), iv) epicuticular synthesis involved P450s (e.g *CYP4G17* and *CYP4G16*), v) transcription factors (e.*g HR38, AP, and ETS21C*), vi) oxidative stress involved genes (including glutathione peroxidase 3, and Nitric oxide synthase) and vii) several unspecified proteins whose functions are not fully annotated (Table S7).

Several detoxification genes potentially linked to insecticide resistance (Fig. S4) were also found to be overexpressed. Some of these genes include cytochrome P450s (*CYP6M4*, *CYP6M3, CYP6M2, CYP9K1, CYP6Z1*, and *CYP4AA1*), UDP-glucosyltransferases (AGAP006223, AGAP007589, AGAP007588) and glutathione s-transferases (*GSTD7*) (Table S7).

#### 3.2.2 Differential expression of genes associated with naphthalene selection

##### 3.2.2.1 Auyo Field Populations

For the field population naphthalene selection (AnNaph), the top commonly overexpressed genes (Table 2) include spinster protein (AGAP006345), Vitamin K-dependent protein C (AGAP009217), Omega-amidase NIT2 (FC = 7.7), ornithine decarboxylase (FC=8.6), unspecified protein (AGAP003757), and Chitin-binding type-2 domain-containing protein (FC=5.6). Commonly overexpressed genes involved in detoxification include CYP4C27 (FC = 3.3), and CYP6M4 (FC=4.1). Other detoxification genes overexpressed in at least one of the comparisons are shown in a heatmap (Fig. S3).

**Table 2:**
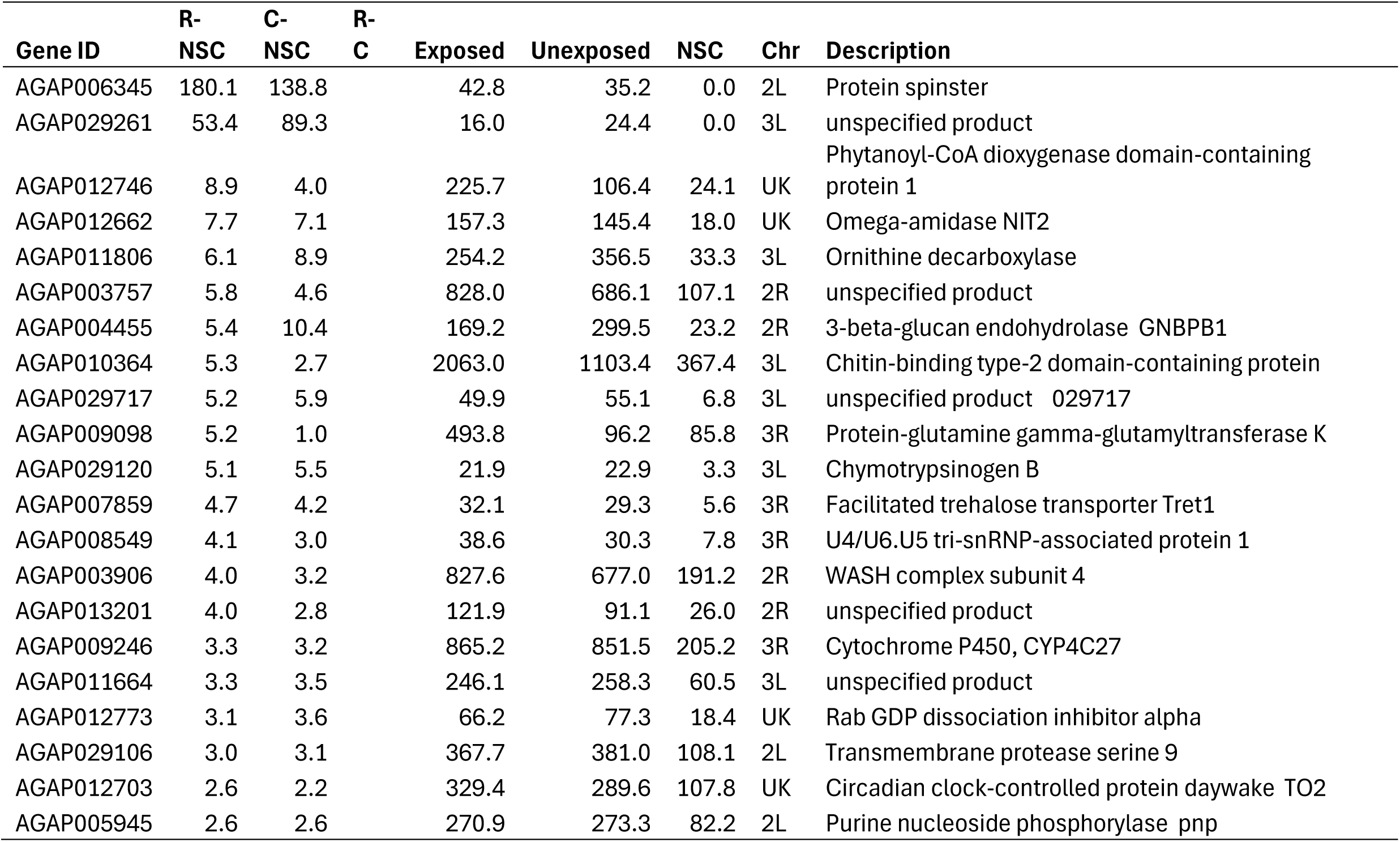
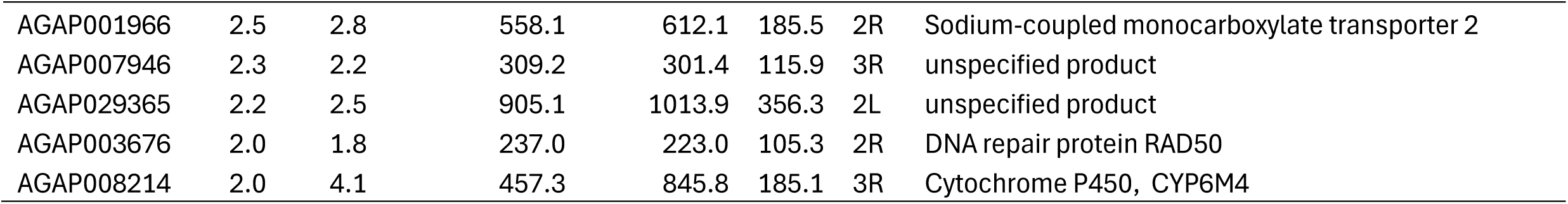
Commonly overexpressed genes in the field population (Auyo) selection with naphthalene (AnNaph) compared to the field non-selected (NS) control. Exposed (R) survived permethrin bioassays at the tenth generation of selection while the unexposed (C) were from the same selected lines but no insecticide exposure (**log_2_FC values >1 and FDR-adjusted *p* < 0.05)**

| Gene ID | R-NSC | C-NSC | R-C | Exposed | Unexposed | NSC | Chr | Description |
| --- | --- | --- | --- | --- | --- | --- | --- | --- |
| AGAP006345 | 180.1 | 138.8 |  | 42.8 | 35.2 | 0.0 | 2L | Protein spinster |
| AGAP029261 | 53.4 | 89.3 |  | 16.0 | 24.4 | 0.0 | 3L | unspecified product |
| AGAP012746 | 8.9 | 4.0 |  | 225.7 | 106.4 | 24.1 | UK | Phytanoyl-CoA dioxygenase domain-containing protein 1 |
| AGAP012662 | 7.7 | 7.1 |  | 157.3 | 145.4 | 18.0 | UK | Omega-amidase NIT2 |
| AGAP011806 | 6.1 | 8.9 |  | 254.2 | 356.5 | 33.3 | 3L | Ornithine decarboxylase |
| AGAP003757 | 5.8 | 4.6 |  | 828.0 | 686.1 | 107.1 | 2R | unspecified product |
| AGAP004455 | 5.4 | 10.4 |  | 169.2 | 299.5 | 23.2 | 2R | 3-beta-glucan endohydrolase GNBPB1 |
| AGAP010364 | 5.3 | 2.7 |  | 2063.0 | 1103.4 | 367.4 | 3L | Chitin-binding type-2 domain-containing protein |
| AGAP029717 | 5.2 | 5.9 |  | 49.9 | 55.1 | 6.8 | 3L | unspecified product 029717 |
| AGAP009098 | 5.2 | 1.0 |  | 493.8 | 96.2 | 85.8 | 3R | Protein-glutamine gamma-glutamyltransferase K |
| AGAP029120 | 5.1 | 5.5 |  | 21.9 | 22.9 | 3.3 | 3L | Chymotrypsinogen B |
| AGAP007859 | 4.7 | 4.2 |  | 32.1 | 29.3 | 5.6 | 3R | Facilitated trehalose transporter Tret1 |
| AGAP008549 | 4.1 | 3.0 |  | 38.6 | 30.3 | 7.8 | 3R | U4/U6.U5 tri-snRNP-associated protein 1 |
| AGAP003906 | 4.0 | 3.2 |  | 827.6 | 677.0 | 191.2 | 2R | WASH complex subunit 4 |
| AGAP013201 | 4.0 | 2.8 |  | 121.9 | 91.1 | 26.0 | 2R | unspecified product |
| AGAP009246 | 3.3 | 3.2 |  | 865.2 | 851.5 | 205.2 | 3R | Cytochrome P450, CYP4C27 |
| AGAP011664 | 3.3 | 3.5 |  | 246.1 | 258.3 | 60.5 | 3L | unspecified product |
| AGAP012773 | 3.1 | 3.6 |  | 66.2 | 77.3 | 18.4 | UK | Rab GDP dissociation inhibitor alpha |
| AGAP029106 | 3.0 | 3.1 |  | 367.7 | 381.0 | 108.1 | 2L | Transmembrane protease serine 9 |
| AGAP012703 | 2.6 | 2.2 |  | 329.4 | 289.6 | 107.8 | UK | Circadian clock-controlled protein daywake TO2 |
| AGAP005945 | 2.6 | 2.6 |  | 270.9 | 273.3 | 82.2 | 2L | Purine nucleoside phosphorylase pnp |

|  |  |  |  |  |  |  |  |
| --- | --- | --- | --- | --- | --- | --- | --- |
| AGAP001966 | 2.5 | 2.8 | 558.1 | 612.1 | 185.5 | 2R | Sodium-coupled monocarboxylate transporter 2 |
| AGAP007946 | 2.3 | 2.2 | 309.2 | 301.4 | 115.9 | 3R | unspecified product |
| AGAP029365 | 2.2 | 2.5 | 905.1 | 1013.9 | 356.3 | 2L | unspecified product |
| AGAP003676 | 2.0 | 1.8 | 237.0 | 223.0 | 105.3 | 2R | DNA repair protein RAD50 |
| AGAP008214 | 2.0 | 4.1 | 457.3 | 845.8 | 185.1 | 3R | Cytochrome P450, CYP6M4 |

##### 3.2.2.2 Ngousso Laboratory susceptible strain

Ngousso selection with showed highly expressed genes, including unspecified protein (AGAP029261, FC= 53), heat shock protein 70kDa (AGAP004581) (FC-= 41), dynein intermediate chain 2, axonemal (AGAP011540) (FC = 26), Protein-glutamine gamma-glutamyltransferase E (AGAP009098) (FC=10), Flavin-containing monooxygenase FMO GS-OX-like 1 (AGAP010398) (FC = 8.5), Phytanoyl-CoA dioxygenase domain-containing protein 1 (FC= 8.9), glucosyl/glucuronosyl transferases (AGAP005754) (FC = 5) and cuticular protein RR-1 family 20 (AGAP005969) (FC = 4). The 70kDa heat shock protein (AGAP004581, FC = 74) was the most overexpressed gene in the selection of the Ngousso line with naphthalene selection (Table S7). Other genes potentially involved in detoxification and insecticide resistance include several cytochrome P450s (including *CYP6Z1, CYP6AA1, CYP6M4*, *CYP6M2)*, GSTs *(e.g GSD7), UGTs (AGAP007588, AGAP007589),* and *ABCs (ABCA2, ABCG1*) (Table S7). Major detoxification genes overexpressed in at least one comparison have been demonstrated in a heatmap (Fig. S4, Additional file 1), and volcano plots of the differentially expressed genes are also presented (Fig. S 8).

#### 3.2.3 Differential expression of genes associated with selection using a mix of fluorene and naphthalene

Selection with the mixture of PAHs (naphthalene and fluorene) led to the overexpression of a combination of genes found in the separate selections with naphthalene and fluorene (Table S6). The selection with a mixture of PAHs may be more realistic to field settings where several PAHs are expected in the environment, and mosquitoes are likely exposed to them simultaneously. The highly overexpressed genes include large ribosomal subunit protein (FC = 46), Histones (including H3.2, H4 and H2A), indicating strong epigenetic marks involved in driving the overexpression of major detoxification genes. Chymotrypsinogen B, ornithin decarboxylase, zinc finger proteins and mitochondrial potassium channel are among the top-expressed genes (Table S6). Some of the commonly overexpressed insecticide-resistance relevant genes include the cytochrome P450s (CYP6M4 and CYP4C27), GSTs (GSTD1-4 and GSTE3). Other detoxification genes overexpressed in at least one comparison are represented in heatmaps (Fig. S 3, additional file 1) while all the differentially expressed genes are shown on volcano plots (Fig. 4 and Fig S8).

##### 3.3.1 Detecting variants/SNPs associated with the generational exposure to PAHs

To understand the impact of PAH exposure on selecting important mutations that are potentially associated with survival to PAHs and/or insecticides, variants were predicted in each line. The scan led to the identification of 5,491 variants while comparing the selected and the non-selected lines in both the field and fully susceptible strains (Fig. S5). Focussing on only missense variants with significant p-values and deep coverage led to the identification of several variants associated with detoxification and cuticle biosynthesis. Furthermore, significant SNPs were found in the cytochrome P450s, GSTs, UGTs, odorant binding proteins and others.

Significant SNPs with low frequencies of up to 10 % in the non-selected controls in the field and laboratory susceptible strain that rose to a higher frequency in the 10^th^ generation of selection were considered to be associated with survival or adaptation to PAHs/insecticides. Such mutations included GSTE4 (Lys190Glu) that rose from 3% to 38% and 49% in the AnMix and AnNaph lines, respectively (Fig. 5c) ; GSTE4- His172Arg with 0% frequency in the control lines but showing up to 27% in the field selection lines was). These two mutations implicate GSTE4 in tolerance/resistance to PAHs and potentially insecticides. Two novel GSTE2 gene mutations (Thr154Ser and Lys92Arg) had frequencies raised from 3% in the control lines to 33% in the field strain selected lines. The cuticular protein CPLCG1 (Thr81Ala) (Fig. 5c) rose from 35% in the Ngousso control to 75% in the fluorene selected Ngousso line (Ngou_Fluo) but with no clear segregation in the field population selection.

**Figure 5:**
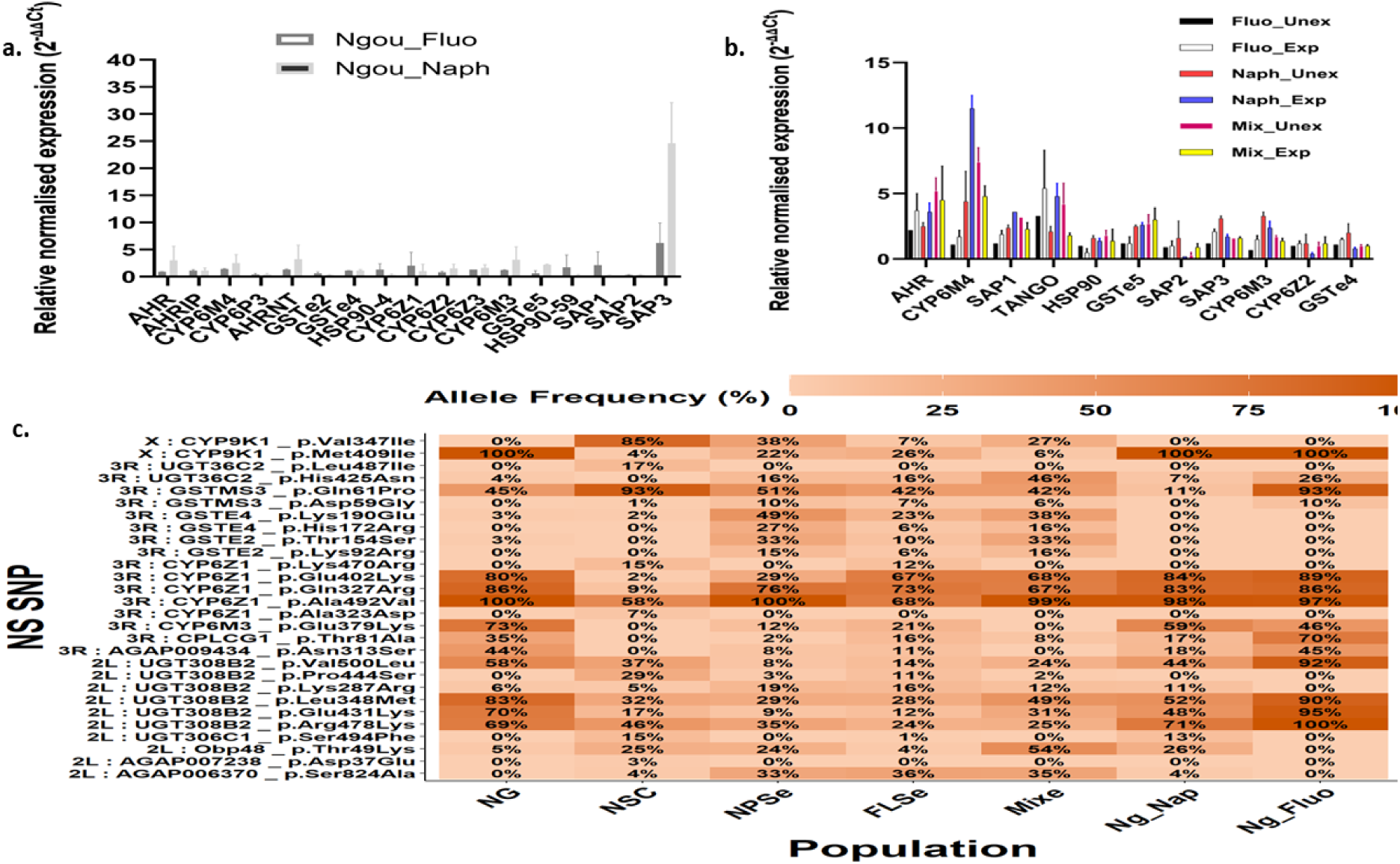
Transcriptional validation of major candidate genes using qPCR and major SNPs detected after selection for 10 generations. The expression profiles of the major genes overexpressed following the selection of the **a**) The laboratory susceptible strains (Ngousso) of *An. coluzzii* and **b**). The field (Auyo). Non-selected lines of the field population were used as a control for the field selection experiments, whereas the Ngousso susceptible strain was used as a control in the Ngousso selection lines. Bars represent the Relative normalised expression (2^-ΔΔCt^) of each candidate gene compared to the house-keeping genes. **c**) Heatmap showing the significant SNPs that are associated with generational exposure of field and laboratory susceptible strains of *An. coluzzii* to PAHs. The populations are represented with shorter names. NG means Ngousso, NSC stands for non-selected control, NPSe stands for the naphthalene selected exposed (AnNaph), FLSe represents the fluorene selected exposed (AnFluo), Mixe represents the AnMix exposed, while Ng-Nap and Ng_Fluo stand for the Ngousso lines selected with naphthalene and fluorene, respectively.

Other clearly segregating significant SNPs which included UGT308B2 (Lys2087Arg) with frequencies raised from 6% in the non-selected lines to 19% and 11% for the AnNaph and Ngou_Nap selected lines respectively. An odorant-binding protein 48 variant (Thr49Lys) increased from 25% frequency in the field non-selected control to 54% following selection with fluorene and naphthalene mixture (Fig. 5c).

##### 3.3.2. Gene-set enrichment analysis of the selected lines of the field and laboratory susceptible populations

Gene ontology and metabolic pathway enrichment analysis were conducted for the significant genes (p <0.05) in the overexpressed genes in the selected versus non-selected lines for the two strains. This showed enrichment of many GO terms and pathways involved in insecticide resistance. For the AnFluo selection, the most enriched terms included transporter activity and transmembrane transporter activity, which contributed up to 28.4% of the overexpressed genes. Odorant-binding genes were also enriched (11.1%). Other enriched GO terms relevant to insecticide metabolism included oxidoreductase activity (8.6%) and monooxygenase activity (6.17%). Some of the notably enriched metabolic pathways in the field population selection on fluorene (AnFluo) include: metabolism of xenobiotics by cytochrome P450, ecdysone and 20-hydroxyecdysone biosynthesis, drug metabolism - other enzymes, drug metabolism - cytochrome P450, and Polycyclic aromatic hydrocarbon degradation pathways. For naphthalene selection, transmembrane transport that included transmembrane transporter activity, transporter activity and secondary active transmembrane transporter activity, was the most enriched GO term, collectively contributing up to 38.4% of the overexpressed genes. Other enriched GO terms linked to xenobiotic/insecticide metabolism include the serine proteases odorant binding and oxidoreductase activity. Metabolic pathways highly enriched in the naphthalene selection include the ecdysone and 20-hydroxyecdysone biosynthesis. The mixed selection (AnMix) appeared to have a higher number of enriched GO terms with strong association with xenobiotic metabolism/insecticide resistance and consistent with the combination of those seen in separate fluorene and naphthalene selections. The most enriched term was organic cyclic compound binding (contributing up to 30% of the expressed genes) involved in the metabolism of cyclic compounds, which potentially include PAHs.

The selection of the laboratory-susceptible strain (Ngousso) with fluorene (Ngou_Fluo) and naphthalene (Ngou_Naph) produced notable enrichment of terms associated with insecticide resistance and metabolic pathways. For example, as for AnFluo selection, Ngou_Fluo produced enriched GO terms that included transmembrane transport genes, which contributed to up to 44% of the total number of overexpressed genes. Ether lipid metabolism pathway involved in oxidative stress and membrane stability enriched in the Ngou-Fluo line along with oxidoreductase activity, serine proteases, and DNA-binding transcription factors.

##### 3.3.3 Transcriptional validation of major candidate genes induced by exposure to PAHs using quantitative real-time PCR

qRT-PCR was conducted on major candidate genes overexpressed in the RNASeq experiment and previously implicated in insecticide resistance or response to environmental xenobiotics. The data supported the RNASeq for most of the genes studied. CYP6M4 generally showed the highest transcriptional induction with fold changes of 1.1 ±0.6, 1.7±0.5 (P = 0.4852), 4.4±2.3 11.5±1.0 (*P* = 0.0473), 7.4±1.1 and 4.8±0.8 (*P*= 0.1273) for the AnFluo, (AnNaph and AnMix, respectively (Fig. 5b). The aryl-hydrocarbon receptor (Ahr) involved in transcriptional regulation of the xenobiotic detoxification genes) (Brown et al., 2005; Mohammed et al., 2016) was also found to be overexpressed with fold changes of up to 5.2±1 in the AnMix selection, followed by 3.7± 1.3 in the case of AnFuo (exposed) for the Ahr and a respective corresponding values of 4.2±1.6, 5.4±2.9 for the AHRNT. The least expressions were recorded for GSTe4, SAP2 and heat shock protein 90 (*HSP90*) (Fig, 5b).

The expression levels of 17 candidate genes were also assayed in the laboratory susceptible Ngousso strain under selection with naphthalene and fluorene. The highest expression levels with fold changes of 24.6±7.5 and 6.2 ±3.7 were recorded for the SAP3 gene (Fig. 5a) (P = 0.0926) in the fluorene and naphthalene selected lines, respectively. *CYP6M4* was also overexpressed with fold changes of 1.4±0.1 and 2.5±1.6 for the fluorene and naphthalene selected lines, respectively (P = 0.5303). For all the genes studied, the naphthalene-selected line (Ngou_Naph) appeared to have higher expression than the fluorene-selected line (Ngou_Fluo). Candidate genes that demonstrated no overexpression in comparison with the non-selected lines include the *GSTe2, CYP6P3, SAP2*, and *GSTe4* (Fig.5a).

##### 3.3.4 Functional characterization of the role of CYP6M4 in the metabolism of pyrethroids and PAHs

In vitro metabolism assays coupled with HPLC analysis revealed that the recombinant CYP6M4 significantly metabolises both types I and II pyrethroids with percentage depletions of 73.8±1.8 %, 72.3±2.3 % and 96.8±0.6% for permethrin, α-cypermethrin and deltamethrin respectively (Fig. 6j). Recombinant CYP6M4 also demonstrated some metabolic activity towards carbamates, propoxur and carbamates with low percentage depletions of 11.2±0.8 % and 17.3 ± 1.5%, respectively. Even though turnover was low, polar metabolites were eluting in both cases, suggesting potential transformation/metabolism of the carbamates. Another important class of compounds tested were the polycyclic aromatic hydrocarbons with representative fluorene, fluoranthene and naphthalene showing percentage depletions of 14.2±2.3 %, 6.9 ±4.4 %, and 16.5 ± 5.6, respectively. Despite low percentage depletions, polar metabolites were visible with retention times of 6.4 min (Fig. 6k) indicating that CYP6M4 metabolises PAHs, but at a much slower rate than pyrethroids. By contrast to these, CYP6M4 showed no activity towards DDT with a percentage depletion of 0.7±1.9%.

**Figure 6:**
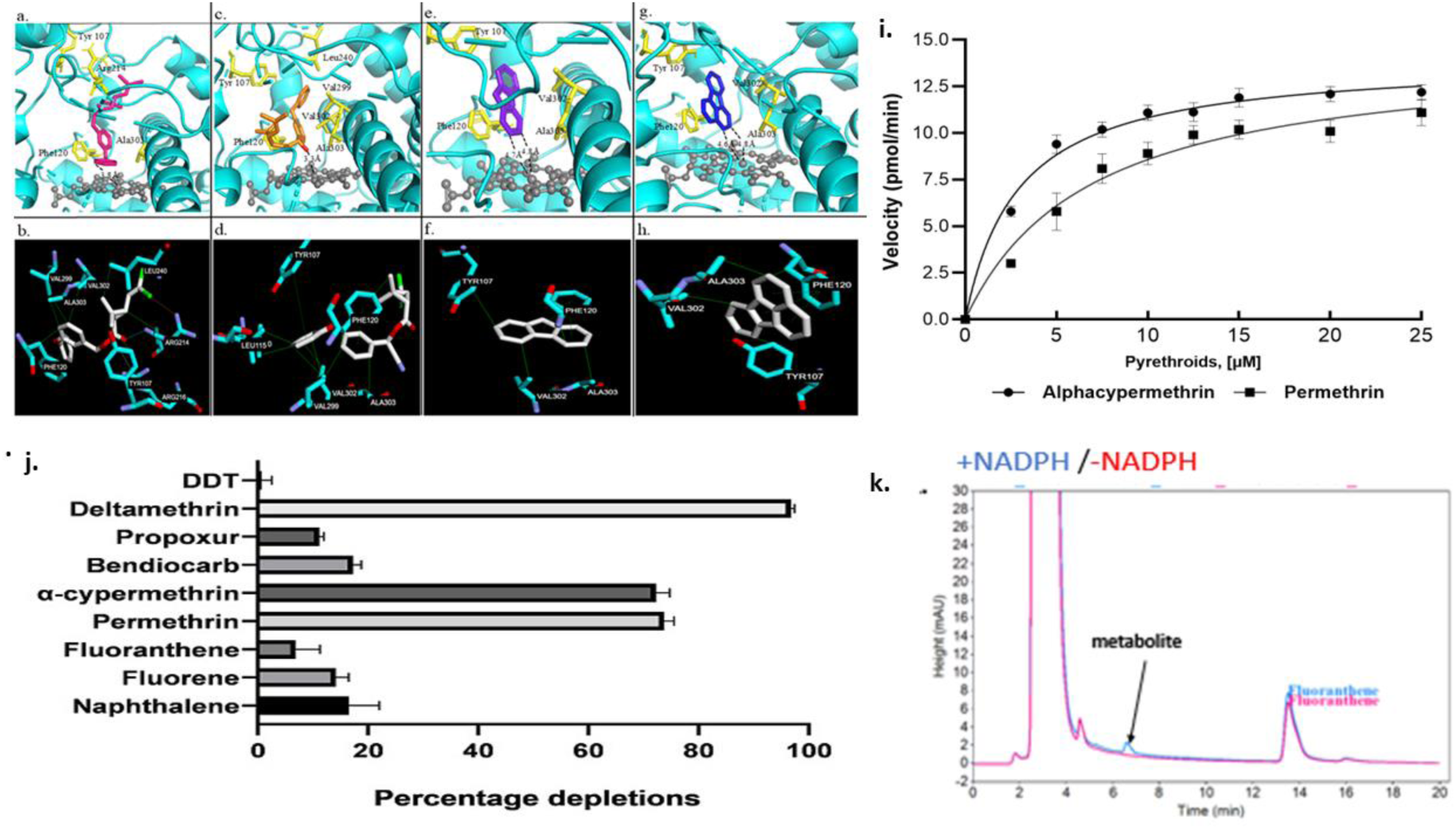
Functional validation of the metabolism of PAHs and pyrethroids by recombinant CYP6M4 using and predicted 3D binding mode of the substrates. Predicted binding mode of (a) permethrin (pink stick), (c) α-cypermethrin (orange), (e) fluorene (purple), and (g) fluoranthene (blue). CYP6M4 helices are presented in cyan colour; heme atoms are in stick format and grey. Distance between possible sites of metabolism and heme iron are annotated in Angstrom. Key residues involved in non-bonded interactions are labelled and in yellow, stick formats. Bottom panels: Key active site residues predicted by protein-ligand profiler, for permethrin **(b)**, α-cypermethrin **(d),** fluorene **(f)** and fluoranthene **(h).** Lime = hydrophobic interactions, cyan = hydrogen bond, blue = water bridge, white = salt bridge, red = *π* cation interaction, green = *π* stacking, orange = halogen bond and grey = metal complexes. **i**) Michaelis-Menten plots of the metabolism of permethrin and α-cypermethrin by recombinant CYP6M4. Values were derived from means of three replicates at each point ± SEM **j**) NADPH-dependent percentage depletions of insecticides and pollutants tested. Values are presented as mean and SD of 3 replicates. **k**) HPLC Chromatogram indicating a metabolite generated from CYP6M4 metabolism of fluoranthene with a 6.4 min retention time.

The turnover rates of permethrin and α-cypermethrin by CYP6M4 were 0.52±0.17 min^-1^ and 0.68±0.12 min^-1^, respectively. The metabolism of permethrin and α-cypermethrin fitted into the canonical Michaelis-Menten model. The steady-state kinetic parameters revealed that while CYP6M4 showed comparable maximal catalytic activity (K_cat_) for α-cypermethrin (0.28 min^-1^) and permethrin (0.29 28 min^-1^), the P450 showed a higher affinity towards α-cypermethrin than permethrin with respective K_m_ values of 2.91 μM and 6.88 μM (Fig. 6i), thus 2.28 x greater catalytic efficiency for the metabolism of α-cypermethrin than permethrin with Kcat/Km values of be 0.10 min^-1^ μM^-1^ and 0.04 min^-1^ μM^-1^, respectively (Fig. 6i).

##### 3.3.4 Prediction of *An. coluzzii* CYP6M4 structurally conserved regions and critical amino acid residues

Alignments of CYP6M4 amino acid sequences to the sequences of previously resolved structures allowed identification of the purported substrate recognition sites (SRSs) and searches using CYPED allowed mapping of the structurally and functionally conserved domains and residues involved in catalysis (Fig. S6). *An. coluzzii* CYP6M4 is 98.2% and 79.6% respectively identical to *An. gambiae* and *An. funestus* CYP6M4 genes. The WxxxR motif, the signatory oxygen-binding pocket (AGFETS)/proton transfer groove, the characteristic ExxR motif which stabilizes the heme structural core, the cysteine pocket/heme-binding region (PFxxGxxxCxG), which forms the fifth axial ligand to the heme iron, and the ‘meander’ (Feyereisen, 2012, Werck-Reichhart and Feyereisen, 2000) were all identical and conserved in *An. coluzzii* and *An. gambiae* sequences, with single amino acid variations observed only in *An. funestus* CYP6M4, with respect to WxxxR motif and the meander. High conservation was also observed in the SRSs, with the only variation between *An. coluzzii* and *An. gambiae* sequences observed in SRS-1 (Asn119Asp).

##### 3.3.5 Prediction of 3D folding pattern of *An. coluzzii CYP6M4* and its ability to metabolise PAHs and pyrethroid insecticides

To investigate the 3D folding pattern of CYP6M4 its homology model was created. Fig. S7, panel a. depicts comparative DOPE energy profiling (internal assessment) between CYP6M4 sequence and the 1TQN (Human CYP3A4) template. Ramachandran energetic validation (panel b) revealed that 377 residues (84.9%), 53 residues (11.9%), and 6 residues (1.4%) were in the most favoured regions, additionally allowed regions, and generously allowed regions, respectively, with only 8 residues (1.8%) in disallowed regions. External assessment using Errat (panel c) provides an overall quality factor score of 46.95% for the best model.

Analysis of the docking parameters and conformation of the ligands in CYP6M4 predicted monooxygenase activity towards both the PAH compounds and the pyrethroids. All four compounds docked with very low binding energies predicted, with hydrophobic interactions contributing greatly in all interactions, while hydrogen bonds were also predicted in the interactions with the pyrethroid insecticides, especially α-cypermethrin (Table S5).

Permethrin docked productively into the active site of CYP6M4 model with the 4′ carbon of the phenoxy ring within 3.8Å from the heme iron, suggesting ring hydroxylation to 4′-hydroxypermethrin (Fig.6a). A total of 8 amino acid residues were involved in non-bonded interactions. Seven of these residues are involved in hydrophobic interactions (Fig. 6b), including Tyr^107^ and Phe^120^, both from SRS-1, involved in interactions respectively, with the cis methyl moiety of cyclopropane ring, and benzyl ring; Val^299^ and Val^302^ and Ala^303^, all aliphatic residues from SRS-4, with the last amino acid being a member of the oxygen-binding pocket; Arg^214^ from SRS-2 vs cyclopropane ring, and Leu^240^ vs the gem trans methyl moiety of the cyclopropane ring. Arg^214^ is also involved in hydrogen bond interaction with the chloromethyl group, as well as a halogen bond to the chlorine atom of the dichlorovinyl moiety.

Despite the strong binding energetics, no hydrogen bonds were predicted for α-cypermethrin; only hydrophobic and *π* stacking interactions were predicted for the type II pyrethroid. α-cypermethrin docked into the active site with the 4′ carbon of the phenoxy carbon ∼11.0Å away from the heme iron (Fig. 6c). However, the benzyl ring oriented to within 4.8Å, suggesting an attack to this aromatic ring. Furthermore, with the α-cyano group ∼3.3Å from the heme iron, oxidative attack to this ester group is also feasible. Residues that involved intermolecular interaction include Leu^115^ and Phe^120^ (both from SRS-2), which are involved in hydrophobic interactions respectively with the phenoxy ring and cis methyl group of cyclopropane ring (Fig. 6d); Leu240 (SRS3) interacting hydrophobically with the phenoxy ring; and there more SRS-4 residues Val^299^, Val^302^ and Ala^303^, all involved in hydrophobic interactions with the phenoxy and/or benzyl ring.

*CYP6M4* was predicted to possess fluorene oxygenase activity. Monooxygenation was predicted through an attack on the 2- and/or 3 position of the aromatic ring to generate dihydrodiols (e.g., 2,3-dihydroxyfluorene) (Fig. 6e). Carbon 2 and 3 are located within 4.7Å and 4.8Å from the heme iron, respectively, with the alicyclic moiety (position 9) of methylenic bridge pointed away at a distance of 8.5Å, suggesting no activity to generate 9-fluorenol. As in the case with the pyrethroids, similar amino acids were predicted to be involved in intermolecular, non-bonded interactions. These include Tyr^107^, Phe^120^, Val^302^ and Ala^303^, predicted to be involved in hydrophobic interactions with C-7, C-4, C-5 and C-3, respectively (Fig. 6f).

Fluoranthene docked into the active site of CYP6M4 model with the C-2 and C-3 oriented above the heme, at distances of 4.8Å and 4.6Å, respectively (Fig. 6g). In this mode, a monooxygenation to produce 2- and/or 3-hydroxyfluoranthene is possible, as well as formation of 2, 3-dione (e.g., trans-2,3-dihydroxyfluoranthene, 2,3-DHFAT). Residues Tyr^107^, Phe^120^, Val^302^ and Ala^303^ were involved in hydrophobic interactions, respectively against C-7, C-2, C-10 and C-2 of the aromatic rings (Fig. 6h).

## 4.1 Discussion

The field population collected from an agricultural site in northern Nigeria used for this selection was found to be highly resistant to pyrethroids and organochlorines, moderately resistant to bendiocarb but susceptible to malathion at F_0_, The exposure of the field strain (Auyo) of *An. coluzzii* to PAHs led to a progressive increase in susceptibility but no significant difference between the selected and the non-selected lines at the tenth generation for permethrin, deltamethrin and DDT but significantly higher mortalities in the control at the 7^th^ generation for deltamethrin and DDT. This may be related to the downregulation of metabolic genes such as CYP6P3, CYP6M2 and CYP9K1 that are generally overexpressed in pyrethroid-resistant populations of *Anopheles*. Thus, mosquitoes with pyrethroid selection pressure removed, the fitness cost associated with the overexpression of metabolic resistance genes is expected to lead to a natural decay in resistance to pyrethroids, as seen in the case of *An. funestus* CYP6P9a-based resistance (Tchouakui et al., 2020). The progressive loss of resistance may also be due to different resistance mechanisms evolving against PAHs against the agricultural origin of the strain. This may be true, especially for the fact that the field population has already been primed for metabolic resistance. In most selection experiments reported earlier (Oliver and Brooke, 2018; Sadia et al., 2024), relatively susceptible or fully susceptible strains were exposed to pollutants, insecticides, and/or pesticides to monitor the changes in resistance over generations. However, in our case, the field strain was already highly resistant from an agricultural background. Future research with a relatively susceptible field strain might give a clearer picture of the mechanism of this selection.

The laboratory susceptible strain selected on PAHs showed reduced susceptibility to pyrethroids and DDT, which is likely because the susceptible strain has lower levels of expression of metabolic resistance genes and fewer/no compounding effects. These observations were in line with previous studies where the selection of *An. gambiae* with agrichemicals led to an increase in deltamethrin resistance (Sadia et al., 2024), while exposure of *Anophelines* to other pollutants/insecticides led to an increase in insecticide resistance levels (Kamdem et al., 2017b; Poupardin, 2014; Sadia et al., 2024). Selections against PAHs led to greater susceptibility to deltamethrin at the tenth generation with both field and laboratory susceptible strains. This could be due to the fitness cost of deltamethrin resistance. It also indicates a possible bioactivation situation between the PAHs and deltamethrin in which generational exposure to PAH increased the efficacy of deltamethrin, which was not observed in the cases of other insecticides tested. Generational exposure of the colony to PAHs might have equally led to the downregulation of deltamethrin-specific metabolic enzymes.

### 4.2 Known and novel metabolic insecticide resistance mechanisms selected/overexpressed in PAHs selection experiments

Whole transcriptome analysis showed constitutive expression/selection of various genes and gene families that have previously been implicated in insecticide resistance, including P450s, which are involved in Phase I oxidative of xenobiotic compounds. CYP6M4 and CYP4C27 were the most consistently overexpressed P450s in field and laboratory strains selections. CYP6M4 was recently found to be downregulated in Sahel populations of *An. coluzzii* (Ibrahim et al., 2023), suggesting its overexpression in Auyo population used in this study was purely due to PAH exposure and thus potentially projected to be involved in PAHs metabolism. However, the overexpression of CYP6M4 has been reported in several other field populations of *Anopheles*, for example, in *An. gambiae* (Wagah et al., 2021) and *An. funestus* (Mugenzi et al., 2023). Despite the consistent overexpression of CYP6M4 in resistant colonies and its implication in insecticide resistance, nothing has been done to functionally validate its potential roles in resistance.

*CYP4C27* has also been implicated in insecticide resistance by overexpression in several field-resistant populations of *Anophelines* (Omoke et al., 2024; Riveron et al., 2017), and in similar selection experiments with insecticides and pollutants (Kamdem et al., 2016). Other P450s overexpressed in at least one of the lines include; CYP6N1, commonly overexpressed in resistant populations of *An. funestus* (Matowo et al., 2022), CYP6P5, a strong metaboliser of pyrethroids in *An. albimanus* (Kusimo et al., 2022), CYP6AA1, a pyrethroid metaboliser (Ibrahim et al., 2018), CYP6P3, a promiscuous substrate metaboliser involved in the resistance to pyrethroids and even carbamates (Kengne-ouafo et al., 2024; Yunta et al., 2019), CYP6Z3 and CYP6Z2 consistently overexpressed in resistant populations (Gueye et al., 2020).

GSTe2, which metabolises DDT, with its overexpression also associated with DDT resistance (Riveron et al., 2014), was overexpressed following laboratory susceptible strain selection with PAHs, with phenotypic resistance to DDT observed after 7 and 10 generations of selection. Another chemosensory gene family highly overexpressed throughout the selection experiments were the odorant-binding proteins known to be involved in olfaction, xenobiotic tolerance and insecticide resistance. Their overexpression may be attributed to the fact that they are involved in the transport and sequestering of hydrophobic molecules like the PAHs and essentially respond to these in a more ligand-receptor interaction pattern (Abendroth et al., 2023). The characteristic smells of the PAHs, especially naphthalene and their hydrophobic structure, are vital grounds for *OBPs* response. Aryl hydrocarbon receptor (Ahr) and its nuclear counterpart, whose roles in the regulation of so many important physiological processes, including response to xenobiotics, were found to be overexpressed (Vogel et al., 2020).

Selection of the colonies on PAHs appeared to also have an impact on the synthesis of the cuticular proteins, suggesting the involvement of the cuticle in tolerating PAHs. *CYP4G16* and *CYP4G17*, involved in the biosynthesis of the cuticular proteins, were found to be overexpressed in some of the lines under selection. This is in addition to the overexpression of several cuticular proteins, including *CPLCP12 (AGAP009759),* and cuticular protein RR-1 family 81*(AGAP011530).* Cuticular proteins are implicated in pyrethroid resistance (Yahouédo et al., 2017) by forming a rigid matrix that reduces the penetration of insecticides to their target sites (Huang et al., 2018). Other gene families with consistent overexpression in the selection experiments were the proteases (serine, cysteine, zinc carboxypeptidases), which differ on the basis of the position or type of proteins they hydrolyse (Ward, 2011).

### 4.3 CYP6M4 overexpressed in the selection experiments is a potent metaboliser of pyrethroids and PAHs

In insects, specifically mosquitoes, pyrethroid metabolism has been shown to proceed primarily via CYP450-mediated hydroxylation at the 4′ carbon of the phenoxy ring (major route), or the *cis*/*trans*-methyl group approaching the heme iron (minor route of hydroxylation) if the 4’ spot is away for optimal interaction to take place (Stevenson et al., 2011). This is true for several P450s from diverse *Anopheles* species, particularly of the CYP6 class of P450s. For example, the closely related *An. gambiae CYP6M2* (Stevenson et al., 2011) metabolises both type I (permethrin) and type II (deltamethrin) pyrethroids (primarily via C-4’ hydroxylation), with lower affinity for permethrin (*K*_M_ = 11 ± 1 μM) compared to *An. coluzzii* CYP6M4 (K_M_ = 6.9 μM). However, *CYP6M2* is also known to metabolise DDT (Mitchell et al., 2012), while CYP6M4 demonstrated no activity towards this class of insecticides. Other P450s which metabolise pyrethroid insecticides via C-4’ hydroxylation include *An. gambiae CYP6P3* which metabolises permethrin (with a depletion of up to 72%) and deltamethrin (Muller et al., 2008). In a related study (Kengne-ouafo et al., 2024), *CYP6P3* metabolised α-cypermethrin with a relatively lower percentage depletion (57.7%±1) and affinity (K_m_ = 7.35 µM ± 2.01) compared to the *An coluzzii* CYP6M4. The duplicated P450s, *CYP6P9a* and *CYP6P9b* from *An. funestus* (Ibrahim et al., 2015) metabolised pyrethroids (permethrin and deltamethrin) with the percentage of permethrin of up to 91.6±2.5 for the Malawi allele of CYP6P9b. These findings were comparable to the depletions seen in the *An. coluzzii* CYP6M4. The same allele showed a depletion of up to 88% of permethrin. CYP6P9a had relatively lower depletions with 65.38±1.5 and 68.38±1.83 for permethrin and deltamethrin, respectively. CYP6M4 had a higher affinity towards permethrin compared to both CYP6P9a/b with respective Km values of 12.68±2.68 μM and 9.9±1.65 μM. CYP6P5 of *An. albimanus* also moderately metabolised permethrin, deltamethrin, and α-cypermethrin with percentage depletions of 30.6% ± 2.8, 56.8% ±1.3 and 57.4% ± 3.8, respectively (Kusimo et al., 2022). These values were all lower than the depletions observed in the *An. coluzzii* CYP6M4. Furthermore, CYP6P5 had a lower affinity (K_M_ = 3.64 μM) towards α-cypermethrin compared to *An. coluzzii* CYP6M4, these may be related to the fact that the metabolism of α-cypermethrin by *An. albimanus* CYP6P5 proceeded via an attack at the C4 prime spot, which is the minor route of metabolism (Kusimo et al., 2022).

Even though the percentage depletions of the carbamates propoxur and bendiocarb were less than 10%, polar metabolites were observed in the HPLC chromatograms, suggesting potential involvement in the metabolism/sequestration of these insecticides at slower rates. These findings were consistent with earlier studies of *An. funestus* CYP6AA1 and CYP6Z1 (Ibrahim et al., 2018, 2016). The metabolism of PAHs by CYP6M4 was also slower than that of pyrethroids, this could be relevant to the delayed toxicity of the PAHs towards mosquito larvae compared to pyrethroids. Moreover, this is very essential for the fact that CYP6M4 overexpression was induced due to PAH exposure, and it eventually metabolized PAHs and pyrethroids.

The metabolism of PAHs has been shown to proceed via C-H activation by cytochrome P450 enzymes in diverse sets of organisms, including humans, bacteria and fungi (Shimada et al., 2018; Srdič et al., 2022). For example, the human *CYP3A4* has been shown using HPLC coupled with NMR to generate a mono-hydroxylated product, fluorenol and a 9-oxo-substituted product, fluorenone, from the parent compound, fluorene (Srdič et al., 2022). Human P450s known to metabolically activate PAHs (e.g., benzo(a)pyrene, naphthalene, phenanthrene, etc) include CYP2A6, CYP2A13, and CYP1B1 (Shimada et al., 2018). A productive pathway for the metabolism of fluorene has been suggested to involve mono- and/or di-oxygenation of C-1, C-2 or at C-3, C-4 (as we observed from docking), with the corresponding *cis-*dihydrodiols undergoing dehydrogenation and then meta-cleavage, which is catalyzed by an extradiol dioxygenase. Oxidation into 3,4-dihydroxyfluorene has been documented to account 80-90% of fluorene by Pseudomonas sp. Strain NCIB9816-4 naphthalene dioxygenase, with 9-hydroxyfluorene (9-fluorenol) accounting for ∼10% of the metabolite (Resnick and Gibson, 1996). This agrees with our findings of CYP6M4 metabolising this PAH and is consistent with our docking prediction. However, alternative pathways of metabolism have been documented.

Fluoranthene degradation is known to proceed via monohydroxylation into monohydroxyfluoranthene, for example, in the bacteria *Sphingomonas sp*. LB126 (Van Herwijnen et al., 2003) and/or dioxygenation at the C-1,2, C-2,3, as well as C-7,8, and C-8,9 in several Mycobacteria species and *Pasteurella* sp. IFA (Rehmann et al., 2001). In humans and rats, fluoranthene is bio-transformed into 1-, 2, 8-phenols, as well as two phenolic dihydrodiols (e.g., 2,3-dihydroxyflyorenthane/2,3-dione), consistent with our prediction of the site of metabolism of his high molecular weight PAH (Day et al., 1992). Several microbes have been described to break down fluoranthene via C-7,8 and/or C-8,9 hydroxylation. For example, soil *Alcalioenes denitrificans* WW1 strain degrades fluoranthene via 9,-10-dihydroxyfluorene metabolism (Weissenfels et al., 1991).

In summary, the transcriptional analysis confirms that *CYP6M4* was induced through PAH selection, while the metabolism assay results and the findings from the docking simulations established it as a cross-resistance gene, capable of metabolising type I and type II pyrethroid insecticides, as well as polyaromatic hydrocarbons, persistent environmental pollutants.

### 4.4 Conclusion

In this study, we characterized the role of the persistent environmental pollutants (PAHs) in the evolution/exacerbation of insecticide resistance through laboratory-based selection experiments, transcriptomics and functional validations. Selection of the field colony on PAHs has shown non-significant changes in insecticide resistance beyond the seventh generation, whereas the laboratory susceptible strain selection showed a significant increase in insecticide resistance. The ubiquitous environmental pollutants PAHs serve as an additional selection pressure driving resistance to pyrethroids through metabolic-based mechanisms, including the overexpression of candidate genes that metabolise the PAHs and pyrethroids. Exposure to PAHs activates an AhR-like xenobiotic sensor in *An. coluzzii,* driving the expression of detoxification genes, increasing survival to pyrethroids. The most important of these includes the CYP6M4 characterised here. The level and nature of pollution in an environment are, therefore, essential factors to consider in designing resistant management strategies or when deploying vector control tools in some of the most polluted urban cities of Sub-Saharan Africa.

## Acknowledgements

The authors would like to acknowledge the Petroleum Technology Development Fund (PTDF) for awarding the overseas scholarship to AM (PTDF/17/OSS/PHD/1113). The project is also supported by Wellcome Trust Senior Research Fellowship in Biomedical Sciences (217188/Z/19/Z) and by the Bill and Melinda Gates Foundation Investment (INV-006003) to CSW. We equally acknowledge the sponsorship of Research Infrastructure for the control of vector-borne diseases: Infravec2, supported by Horizon 2020 (Grant Agreement No. 731060) for conducting the RNASeq in their laboratory.

## Data Accessibility and Benefit-Sharing

The dataset(s) supporting the conclusions of this article have been deposited under the project code PRJEB106951 in the European Nucleotide Archive (ENA). The cDNA sequence of *CYP6M4* has been deposited in GenBank and the accession number (PX866997) has been obtained.

## Author’s contribution

AM, CSW, MJIP and SSI conceived the idea and designed the experiments; AM carried out the laboratory investigations. HI, TA, AT, and JH, were involved in data analysis and curation. CSW and MJIP provided the oversight/supervision and data validation. AM and SSI came up with the first draft of the manuscript. All authors have read and approved the last version of the manuscript

